# Reverse Engineering DNA Origami Nanostructure Designs from Raw Scaffold and Staple Sequence Lists

**DOI:** 10.1101/2023.05.03.539261

**Authors:** Ben Shirt-Ediss, Jordan Connolly, Juan Elezgaray, Emanuela Torelli, Silvia Adriana Navarro, Jaume Bacardit, Natalio Krasnogor

**Affiliations:** Interdisciplinary Computing and Complex Biosystems Research Group, School of Computing, Newcastle University, Newcastle-upon-Tyne, NE4 5TG, UK; Centre de Recherche Paul Pascal, CNRS, UMR5031, 33600 Pessac, France

## Abstract

Designs for scaffolded DNA origami nanostructures are commonly and minimally published as the list of DNA staple and scaffold sequences required. In nearly all cases, high-level editable design files (e.g. caDNAno) which generated the low-level sequences are not made available. This de facto ‘raw sequence’ exchange format allows published origami designs to be re-attempted in the laboratory by other groups, but effectively stops designs from being significantly modified or re-purposed for new future applications. To make the raw sequence exchange format more accessible to further design and engineering, in this work we propose the first algorithmic solution to the inverse problem of converting staple/scaffold sequences back to a ‘guide schematic’ resembling the original origami schematic. The guide schematic can be used to aid the manual re-input of an origami into a CAD tool like caDNAno, hence recovering a high-level editable design file. Creation of a guide schematic can also be used to double check that a list of staple strand sequences does not have errors and indeed does assemble into a desired origami nanostructure prior to costly laboratory experimentation. We tested our reverse algorithm on 36 diverse origami designs from the literature and found that 29 origamis (81%) had a good quality guide schematic recovered from raw sequences. Our software is made available at https://revnano.readthedocs.io.

## I. INTRODUCTION

DNA origami nanostructures^1,2^ are finding increasing application in diverse areas of nanobiotechnology, such as in single molecule biosensing^3^, targeted drug delivery^4,5^, cell gene delivery^6^, enzyme cascade engineering^7^, nano-patterning^8^ and molecular robotics^9^.

While recent efforts are attempting to define a universal nanostructure exchange format^10^ or to set up a centralised database to store electronic origami design files^11^, origami nanostructure designs are traditionally exchanged in their most minimal form: as raw staple and scaffold sequence lists in publication supplementary material. In a few cases, sequence lists are accompanied by a static low-resolution picture of the origami schematic (e.g. caDNAno^12^ diagram), often omitting sequence information and base pair locations. Only very rarely are editable electronic source schematics made available at publication.

This state of affairs means that published origami nanostructures may be attempted in the laboratory again (in an unchanged form) by third parties, but it is not possible to examine a structure in more detail than is initially provided, and it is difficult to significantly tweak, modify or re-purpose a published design for new future applications.

All current computational CAD tools for DNA origami^1,13–17^ focus on the forwards problem of creating a nanostructure design and then deriving the staple sequences which realise it (Figure 1a). Physical self-assembly under appropriate reaction conditions in the lab then ‘molecularly’ solves the inverse problem of assembling the staple and scaffold strands back into the designed origami nanostructure. However, the actual self-assembly reaction is an extremely complicated and ill-understood ‘black box’ process^18,19^, and raw strand sequences mean little to human origami designers.

**FIG. 1.**
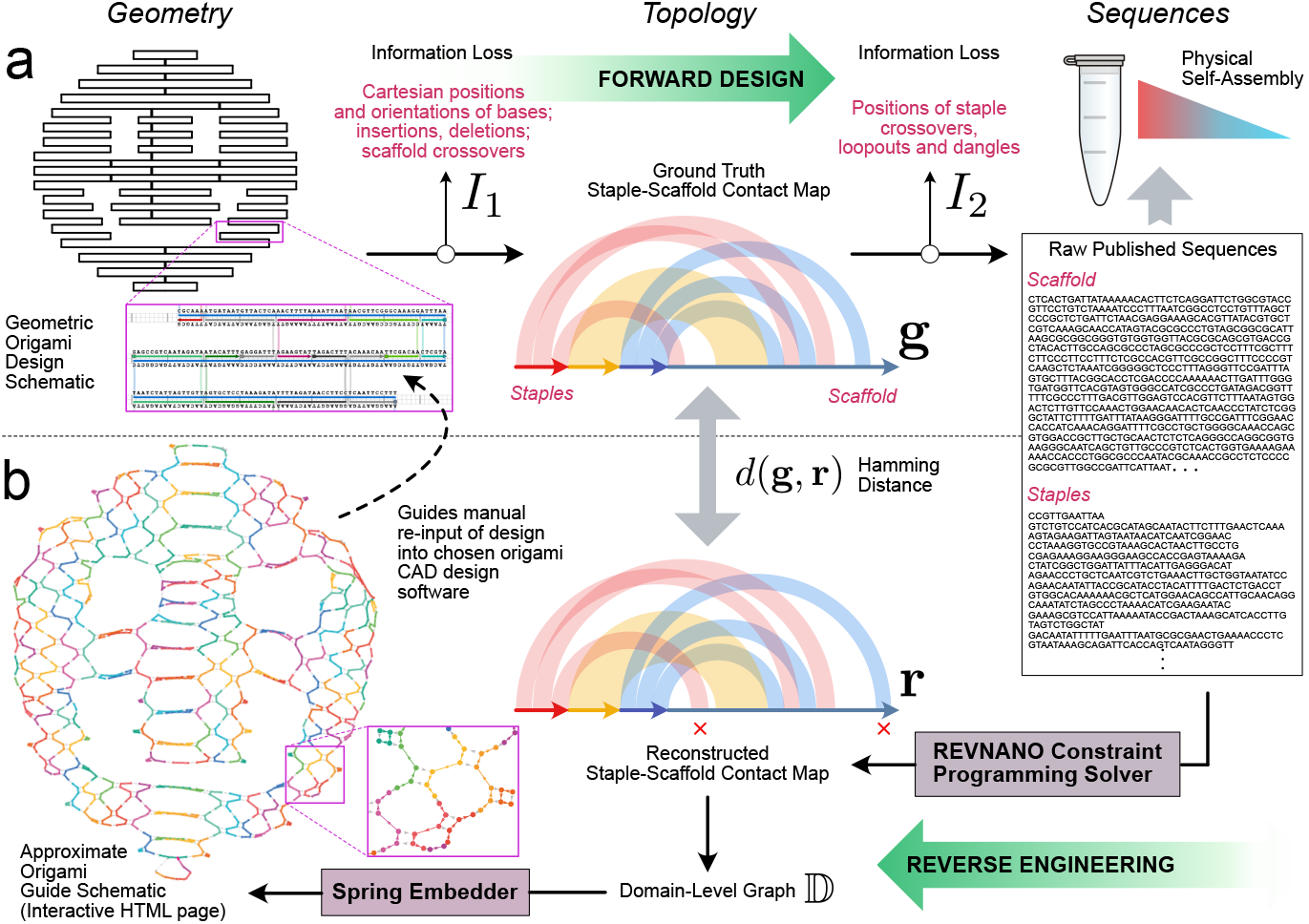
Forward Design and Reverse Engineering of DNA Origami Nanostructures. **(a)** In the traditional forward design process, a DNA origami schematic has a scaffold sequence assigned from which the Watson-Crick complementary staple strands are derived. The geometric origami schematic implicitly embeds a topological contact map detailing how the staple and scaffold bases are pairwise hybridised. The process of going from schematic to raw sequences involves two stages of information loss, *I*_1_ and *I*_2_. **(b)** The inverse problem addressed in this work, i.e. going from raw sequences back to a geometric origami schematic, involves the (partial) recovery of *I*_2_ and then *I*_1_. The REVNANO constraint programming solver first reconstructs an approximate contact map of scaffold-staple base pair connectivity **r**; the latter is converted into an equivalent graph representation of the origami domain-level connectivity D; finally a spring-embedder algorithm converts the graph into an approximate non-crossing geometric representation in 2D or 3D using spring energy minimisation.

To make origami designs published as raw sequence lists more accessible to further design and engineering, in this work we propose the first algorithmic solution to the inverse problem of converting sequence lists back to a ‘guide schematic’ resembling the original origami design schematic (Figure 1b). The guide schematic is an interactive HTML page in 2D or 3D, showing basic helix connectivity, staple crossovers and sequence positions on the origami. Rather than attempting to simulate the complex physical self-assembly pathway of scaffold and staple strand hybridisation, our reverse engineering approach instead reconstructs an origami guide schematic in a direct way using staple-scaffold sequence complementarity, constraint programming and graph layout algorithms. These techniques allow to partially recover information lost in the forward origami design process (marked as stages *I*_1_ and *I*_2_ in Figure 1a).

Reverse engineering an origami guide schematic from raw published sequences is useful for a number of reasons. Firstly, the guide schematic can be used to aid the manual re-input of the origami nanostructure into an origami CAD design tool of choice, hence recovering an editable electronic schematic which can be modified and extended for future research. During this process, images published of the original schematic can also be used as a complementary information source.

Secondly, the guide schematic allows to see in greater detail origami design features that were not mentioned, or which were described only vaguely in the original origami publication; for example, whether an origami possesses a loop of excess single strand scaffold and how staples are hybridised within the loop.

Thirdly, the existence of a reverse engineered guide schematic provides validation that a particular set of staple sequences do indeed lead to an intended origami shape, i.e. no strands are missing and no strands have sequence errors. This is useful both for authors publishing sequences (to verify that no typos or omissions are made) and for third parties wanting to replicate published designs (to verify listed sequences are correct before going to the expense of fabrication).

Fourthly, if a publication is not explicit about the origami scaffold sequence used (a surprisingly common occurrence), then the staple sequence set can be reversed against different scaffold sequences until the correct one is found.

Finally, the ease of reverse engineering an origami design from sequences is likely roughly correlated with decreased likelihood of kinetic traps during origami self-assembly, as staple addressability is improved (see Discussion).

Our reverse engineering pipeline first recovers a staple-scaffold contact map of an origami design, detailing which staple and scaffold bases are pairwise hybridised (see Supplementary Note 1). Construction of contact maps from raw sequences mined from publications could also, in its own right, underpin future data science / machine learning applications in structural DNA nanotechnology (see Discussion).

It should be noted that simple counterexamples demonstrate that the origami reverse engineering problem is not solvable in the full general case. For example, if an origami design has a pathological scaffold sequence applied, like a single-letter sequence (e.g. polyA) or a repeating sequence (e.g. ATATATAT…), then the complementary staple sequences derived will be too homogeneous to contain sufficient information to reconstruct the origami schematic. However, the full general case is not of practical interest as the latter pathological sequences will likely not self-assemble properly either. In practice, origami sequences do contain enough local sequence variety to ascertain the original locations of staples by sequence complementarity. Hence, in practice, the inverse problem can be attempted.

Also, it should be noted that the reverse engineering problem does not admit a trivial solution. For instance, it may seem that the following algorithm (to be named NAIVE) would be sufficient to reconstruct the staple-scaffold contact map for an origami, from just sequence information: 1) Run a staple strand anti-parallel to the scaffold strand; 2) Record all regions where there is a complementary staple-scaffold sequence match from the 5’ of the staple, over a certain minimum number of base pairs *μ*_min_; 3) Take the longest of these regions as the route of the staple; 4) Truncate the staple to not include the matched region; 5) Repeat from (1) until all bases of the staple strand are matched; 6) Repeat for all staples.

However, while NAIVE does work with some origamis (see Results) it fails on other designs, such as origamis based on the M13 scaffold with a high number of repeated sequence regions (which can, in some instances, entirely replicate an intended staple binding site).

In physical self-assembly, staples eventually find their correct hybridisation locations on the origami scaffold strand due to cooperative loop formation and cascades of strand displacement reactions that constantly re-arrange binding configurations over a gradual temperature annealing ramp until myriad off-target bindings are statistically cleared away. The view that each staple binds only at one set of designed locations, as assumed by NAIVE, is often too simplistic.

In this work, we develop a two-stage pipeline (Figure 1b) to robustly solve the inverse problem of raw staple/scaffold sequences to origami guide schematic. Described in more detail below, in the first stage of this pipeline we develop a constraint programming solver called REVNANO to recover the (approximate) staple-scaffold contact map from origami sequences. In the second stage of the pipeline, we use graph layout techniques to convert the topological contact map into an approximate geometric origami schematic. In the Results section we describe good reverse engineering performance of our algorithms on a diverse test set of 36 origami nanostructures drawn from the literature, and analyse failure cases in detail. In the Discussion we review current limitations of our approach and highlight future improvements which could be made to our work.

## II. REVERSE ENGINEERING PROCEDURE

The problem of reverse engineering a DNA origami guide schematic from raw scaffold and staple sequences can be split into two independent sub-problems, each described below.

### A. Recovery of Origami Contact Map from Sequences

The first sub-problem is to recover a good approximation of the contact map for the origami design, given just the raw scaffold and staple sequences. The contact map simply states which scaffold and staple bases are hybridised together, and which are left unpaired in the origami design (see Supplementary Note 1). With no notion of spatial positioning, a contact map is a purely topological representation of base pair connectivity within a folded origami. During the forward design process (Figure 1a), the contact map for an origami nanostructure is actually embedded implicitly in the geometric schematic. However, in the reverse process, we are required to recover the topological contact map explicitly.

Contact map recovery falls into the class of constraint satisfaction problems^20^ (CSP) in Computer Science. In a CSP, a set of decision variables exist where each variable may assume one of a finite set of alternative values. The problem is to find an assignment of values to the decision variables such that a particular set of constraints are satisfied on individual- and/or between groups of decision variables. For the origami case, the decision variables are staples. The set of alternative values for each staple are the set of potential Watson-Crick complementary routes through the scaffold that each respective staple may assume. The problem is to assign one route to each staple such that (i) all staples are routed on the scaffold and (ii) no staple-staple overlaps exist. It will generally be the case that only a single unique solution exists to this problem, due to high staple packing density on a DNA origami covering most or all of the scaffold strand.

We implemented a custom constraint programming solver called REVNANO to solve the CSP leading from raw scaffold/staple sequences to the origami contact map. The solver leverages the unique physical features of origami nanostructures as heuristics to arrive at a solution faster. DNA, RNA^21,22^ or hybrid scaffolded origami^23,24^ are all supported. The core principles of the solver are outlined below and exemplified in Figure 2 using the Rothemund smiley face origami^25^.

**FIG. 2.**
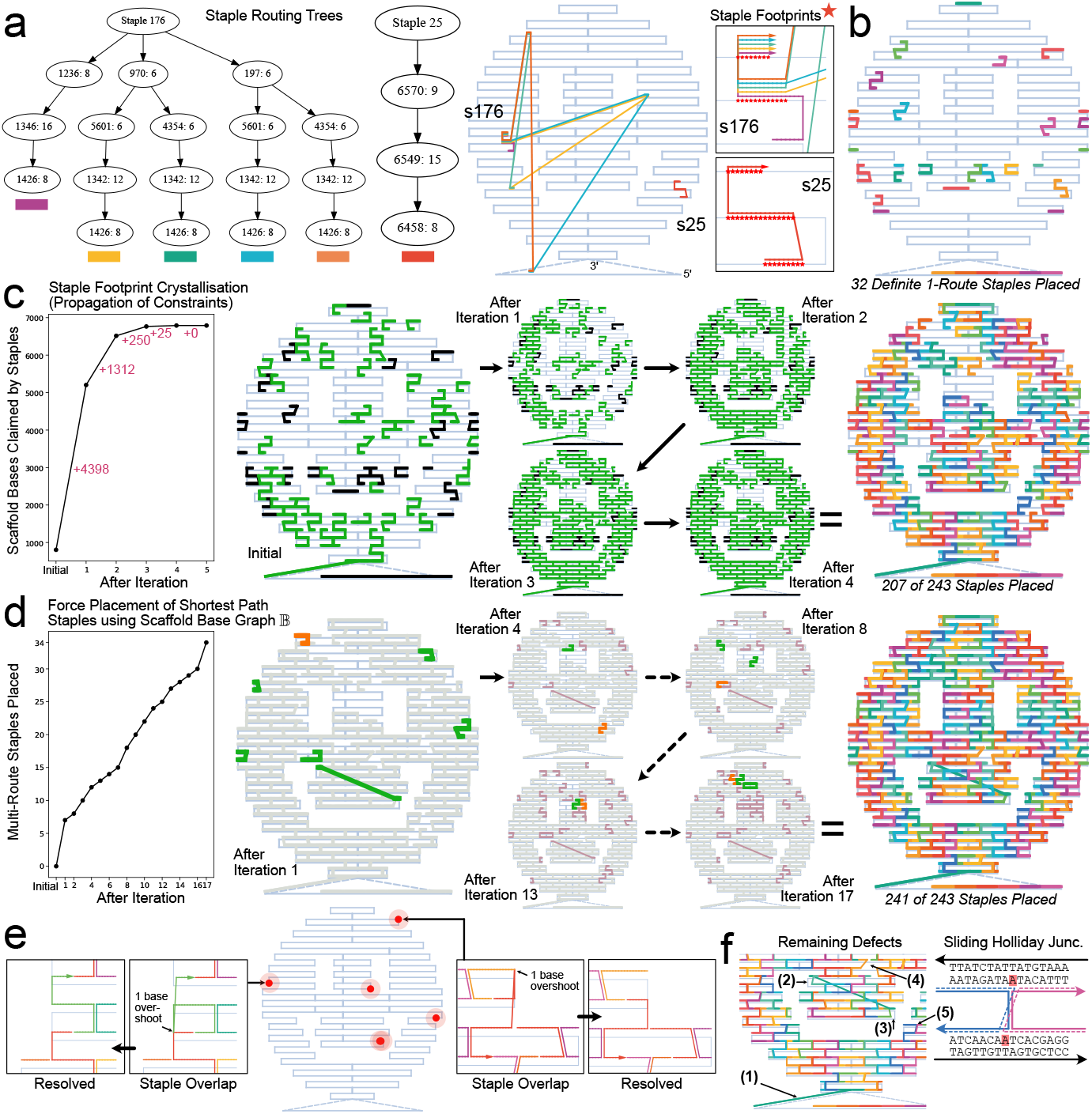
REVNANO Constraint Programming Solver: Principles Illustrated with Rothemund Smiley Origami^25^. **(a)** Stage 0: Staple routing trees (SRTs) are constructed for all staples. Tree node (1236: 8) signifies a staple section that begins at scaffold base 1236 and runs for 8 bases in total toward scaffold 5’. Staple 176 has 5 viable routes across the scaffold; Staple 25 has 1 single viable route and casts a larger ‘footprint’ (inset boxes, red starred bases). **(b)** Definite 1-route staples are identified and placed on the origami as initial hard constraints. **(c)** Stage 1: SRTs are pruned by the footprints of other SRTs in an iterative process. Over successive iterations more staples (green) collapse to a single defined route. The initial hard constraint staples placed at Stage 0 are shown in black. **(d)** Stage 2: ‘Problem’ staples remaining with >1 route are placed by forcing the staple with the clearest shortest path through the current origami mesh to its shortest path route, and repeating over and over. Orange = staple forced to single route on iteration; Green = ‘ripple effect’ staples collapsing to a single defined route because of the latter action; Purple = total problem multi-route staples placed up to the current iteration. **(e)** Stage 3: Staple-staple overlaps (at locations shown by red dots) are fixed. **(f)** A few minor defects remain (see Results for discussion). For the REVNANO parameters used in this example (*μ*_min_ = 6bp, *σ* = 5bp, *β* = 0.5), 240 staples are placed approximately correctly, 1 staple is placed incorrectly and 2 staples are omitted. Note that in all diagrams, staples are superimposed on top of a pre-existing smiley scaffold routing to clearly show how staples have been placed: however, REVNANO does not know this scaffold routing *a priori*.

At Stage 0 of the REVNANO solver, a staple routing tree (SRT) is constructed for each staple. In a staple routing tree, nodes represent scaffold regions where individual sections of a particular staple hybridise, and edges represent staple crossovers (Figure 2a). The root node of the SRT represents staple 5’, and tree branches represent different staple routes through the scaffold (toward staple 3’). The SRT of each staple is computed independently of the other staples.

Two parameters are important in SRT construction. Parameter *μ*_min_ is the minimum length complementary region between staple and scaffold that is permitted to be a routing tree node. In practice *μ*_min_ = 5bp or 6bp. Conversely, parameter *σ* controls the number of alternative competitor sibling nodes allowed in each subtree and makes SRT size computationally tractable. To give an example, if a child node in a particular subtree represents a match of *μ* = 16bp between staple and scaffold, and this is the highest of its siblings in that subtree, then when *σ* = 5, other sibling nodes must represent a match of at least *μ* = 16 - 5 = 11bp in order to also be included in the subtree. In practice, 0bp ≤ *σ* ≤ 10bp.

For the smiley face origami in Figure 2, 34 staples have an SRT with exactly one route (like Staple 25, Figure 2a) and 209 staples have an SRT with more than 1 route (like Staple 176). The latter multi-route staples have, on average, 60.2 possible scaffold routes per staple and one staple has a staggering 1208 possible scaffold routes.

Each staple routing tree casts a ‘footprint’ on the scaffold (red stars in inset boxes, Figure 2a), which is defined as the set of scaffold bases that all routes in the SRT pass through: in other words, the set of scaffold bases definitely claimed by the staple. Footprint size increases as the number of potential scaffold routes a staple has decreases, and reaches a maximum when only 1 viable route exists.

Stage 0 finishes by placing definite 1-route staples on the scaffold (Figure 2b).

In Stage 1 of the REVNANO solver, constraint propagation happens. The full footprint (true route) of each staple gradually ‘crystallises out’ over a series of iterations (Figure 2c). In the first iteration, staples are polled either in order (deterministic mode, the default used in this paper) or at random (non-deterministic mode, see Supplementary Note 4). Each staple prunes its routing tree in response to the current footprints cast by the other staples. Some branches in the SRT of a polled staple will no longer be viable as they intersect scaffold bases already claimed by other staples. When the SRT of a polled staple is pruned, the staple casts an enlarged footprint on the scaffold. After all staples have been polled once, iteration 2 starts and all staples are polled again. The process iterates until no staple enlarges its footprint (typically 5 - 10 iterations are required). A third REVNANO parameter is relevant at this stage: parameter 0 *< β* ≤ 1 controls how much overlap of staples is tolerated during staple placement. If a staple route wants to claim a certain region of the scaffold that has some portion already claimed by another staple footprint, then that region is unavailable if *β* fraction (or greater) of its bases are already claimed, otherwise it is available. Parameter *β* relaxes the requirement that staples should be strictly non-overlapping in the early stages of the solver. Initial fuzzy staple boundaries are often essential to obtain a good final solution in many cases.

Constraint propagation during Stage 1 typically reduces the majority of staples to a single route. However, a small group of staples often resists this purely deductive approach. This ‘G1’ group of staples remain with multiple potential routes. In the Figure 2 smiley example, the G1 group is 15% of staples. At this stage a standard CSP solver may use forward- and back-tracking to solve the remaining decision variables. However, REVNANO leverages the fact that origami nanostructures are physical, highly connected objects to reach a solution more efficiently.

At Stage 2 of the REVNANO solver, a scaffold base graph 𝔹 is constructed using the already placed staples. In the graph 𝔹, two scaffold bases (nodes) are linked by an edge if a staple crossover exists between them, or if the scaffold backbone connects them. Because the majority of staples are placed at Stage 1, the scaffold base graph is a highly connected mesh and shortest path distances between any two scaffold bases well-approximate true shortest path distances in the hypothetical base graph of the completed origami. REVNANO fails with an error message at Stage 2 if less than 65% of staples are placed in Stage 1 (independent of REVNANO parameter values).

Stage 2 of the REVNANO solver uses the scaffold base graph 𝔹 with each G1 staple in the following way: it assigns a shortest path value (i.e. the least number of bases traversed through the scaffold base graph, from scaffold base where staple 5’ hybridises to scaffold base where staple 3’ hybridises) to each route in the G1 staple SRT. The G1 staple with the ‘clearest shortest path’ property is forced to reduce to its shortest path route, enlarging its staple footprint (orange staples, Figure 2d). The staple with ‘clearest shortest path’ property is defined as the staple with the greatest distance between its shortest path route and its next-shortest path route (i.e. the route when the shortest path route is prohibited). The clearest shortest path staple is the staple most likely to have its shortest path route correct due to the large margin to the next alternative routing path for the staple. If two or more staples share the clearest shortest path property, the staple with the highest ID number is selected.

Force placement of the clearest shortest path staple generally has a ripple cascade effect where other staples then also collapse to a single route (green staples, Figure 2d). Once the ripple cascade effect has died out, the scaffold base graph B is updated with the newly placed staples, and the next iteration begins. The staple with clearest shortest path in the remaining G1 staple set is identified and again force-placed. Iterations continue until either all G1 staples are solved or no G1 staple with a clearest shortest path can be identified any more. Staples with more than 1 possible route after Stage 2 are omitted from the final reverse engineered origami contact map.

The use of (i) a well-connected origami base graph 𝔹 and (ii) the concept of clearest shortest path, represents a very conservative heuristic to “guess” the correct locations of multiroute staples by using physical distances. This dispenses with the need for forward- and back-tracking at this stage of the CSP solver.

Stage 3 of the REVNANO solver corrects any staple-staple overlaps by remedying overshoots of staple sections. The smiley example in Figure 2e has five such overlap sites.

Note that staples with end dangles and interior loop-out regions that do no hybridise the scaffold (e.g. polyT ssDNA regions) must have these subsequences pre-marked in the sequence input file (see Supplementary Note 6 for a discussion of this important issue). Before running, the solver splits these staples into a series of smaller ‘virtual’ staples, with each virtual staple being one hybridising region of the original staple. Stage 4 of the solver re-assembles these virtual staples into real physical staples once the hybridising parts have been routed. Automatically identifying non-hybridising sections of staples turns out to be a computationally complex and likely intractable problem.

The REVNANO solver only uses sequence matching as constraints on the routing of staples. DNA helix twist may appear to be another type of geometric constraint limiting the positions of staple crossovers. However, helix twist is not leveraged, since many origami designs have intentional base insertions and base deletions at selected points in order to minimise nanostructure bending and twisting^26^. Such insertions and deletions mean that helix twist does not stay uniform at 10.5bp per helix turn (for DNA) on all parts of the origami. Conversely, staple-scaffold complementary is universally respected across all origami nanostructures, whether or not they have base insertions/deletions and regardless of whether the material is DNA, RNA or hybrid.

When an approximation of the origami contact map has been solved, the quality of the REVNANO solution is quantified by taking the base hamming distance between the ground truth contact map **g** for the origami shape and the contact map recovered from sequences **r**. The base hamming distance *d*(**g, r**) is simply the number of scaffold bases in **r** which must be substituted (i.e. have an incorrect hybridisation partner) in order to arrive at the correct solution in **g**. When hamming distance is 0, all staples are placed perfectly. Note that REVNANO does not have access to **g** when constructing the contact map **r**. See Supplementary Note 2 for how the ground truth contact map for each origami nanostructure was derived.

To contrast with the REVNANO solver, we also implemented the NAIVE solver described in the introduction (which replaces REVNANO Stages 0,1 and 2). The NAIVE solver can be seen as solving the degenerate case of the CSP problem where each staple only has one single possible route, and thus each staple can be solved in isolation of the others. The NAIVE solver has one single free parameter *μ*_min_.

### B. Recovery of Geometric Guide Schematic from Origami Contact Map

After an approximate contact map has been solved, the second sub-problem is to turn the base-pair connectivity description of the origami into a geometric layout (guide schematic) which embeds that connectivity into 2D or 3D Cartesian space.

Previous studies have described the reconstruction of the 3D solution shape of DNA origami from design schematics by using finite-element methods to solve the ground state mechanical structure iteratively (CanDo^27–29^) or by using coarse-grain simulation (e.g. MrDNA^30^ or oxDNA^31,32^). To do so, these works contain detailed mechanical descriptions of ss-DNA, dsDNA and different junction types. We note that the aim in the current work is quite different: to recover an interactive representation of helix connectivity in an origami design, which clearly marks out staple/scaffold crossover positions and sequences. While the final geometric layout should approximate the final shape of the origami, mechanical accuracy is not a prime concern.

To approach the problem, we first convert the origami contact map into a domain-level graph 𝔻, similar to that proposed in Ref^33^. In 𝔻, graph edges have five types: single-stranded scaffold domain, double-stranded scaffold domain (where a staple section is hybridised), scaffold nick, staple crossover (standard crossover, or staple loop-out) and staple dangle. Nodes of the domain-level graph are base positions on the scaffold where the above domains start and finish. The graph 𝔻 has a 1:1 mapping with the origami contact map. See Supplementary Note 1 for more information on the domain-level graph.

The domain-level graph 𝔻 is next converted to a geometric layout (i.e. a Cartesian coordinate is derived for each graph node) by the Kamada-Kawai spring embedder^34^, a generic graph layout algorithm. In essence, the Kamada-Kawai algorithm sets up a graph as a mechanical system where graph nodes are stiff rings connected by springs. The algorithm computes the shortest path between all graph node pairs (computes an all pairs shortest path) and uses these graph-theoretic distances as proxy for the desired Euclidean distance between node pairs. Nodes with the longest graph route separating them are considered to be furthest away in Cartesian space. The spring system is set up in such a way that minimising the energy of the system corresponds to all node pairs separated by a Euclidean distance equal to their shortest path distance.

In general, Kamada-Kawai tends to produce a good layout for origami schematics, since the direct Euclidean distance between two points on an origami schematic is generally highly correlated with the Manhattan distance through the scaffold and staple crossovers linking the two points. The use of whole scaffold domains as edges in graph 𝔻 reduces the complexity of the layout task (as opposed to representing each base individually and representing both strands of each helix) and also ensures that DNA helices are displayed as straight lines in the geometric guide schematic. In particular, we found the Python implementation `networkx.drawing.layout.kamada_kawai_layout()` to be a good candidate for producing robust geometric layouts in both 2- and 3-dimensions, starting from a circular configuration of graph nodes.

The geometric layout of graph 𝔻 is then put into a further graph-layout physics engine^35,36^ where small repulsive forces are defined between graph nodes. This latter engine helps to resolve any remaining graph edge crossing, and allows interactive translation, rotation and zooming of the graph on an HTML page. It also provides a reactive physics-based response to node dragging which can be used to manually disentangle parts of a design.

Finally, we developed a general algorithm AMBIG which uses the graph 𝔻 to quantify by how much each Holliday junction or half-crossover junction on the origami can move and still retain sequence complementarity with the scaffold. On the guide schematic, AMBIG explicitly highlights which junctions are ambiguous and may not be properly placed by the solver, and which junctions are non-ambiguous, immovable and are hence correctly placed. If all junctions are immovable then information loss *I*_2_ = 0 in Figure 1a.

## III. RESULTS

### A. REVNANO Performance

We first evaluated the performance of the REVNANO solver over a heterogeneous test set of 36 origamis, comprising 17 raster origamis made on hexagonal or square grids in caDNAno^12^ or scadnano^15^, and 18 wireframe origamis routed by the ATHENA software^37^. Overall, 25 origami designs were 2-dimensional and 11 designs were 3-dimensional. All origamis tested were DNA origami, except for one hybrid origami with RNA scaffold and DNA staples (Origami 1).

Table I shows that, under optimal parameter settings, REVNANO could reverse engineer an approximate contact map for 94% of origamis tested. Further, for 81% of origamis, over 95% of staples could be routed and for 47% of origamis, all 100% of staples could be routed. Worse case performance was observed for Origami 9 (Membrane Nanopore) with 83.7% of staples routed. As expected, no significant differences existed when reverse engineering contact maps for 2D or 3D designs.

**TABLE I.** REVNANO Solver Performance on 36 Origami Test Set. Performance of the NAIVE solver is listed for comparison. For each origami, both solvers are used at their optimal parameter settings (listed). Origami 10 is similar to – but not exactly the same as – the single staple origamis proposed in Ref^44^. Wireframe shapes (apart from Origami 33) were generated using default examples in the ATHENA software^37^ (with 42bp edges). Sequences used were those output by the ATHENA software. For the other origamis, the published staple and scaffold sequences were used. Shorthand: Dim = 2D or 3D origami; *d*(**g, r**) = Base hamming distance from ground truth contact map **g** to reconstructed contact map **r** for the origami shape; SP% = percent staples placed (but not all staples may be routed correctly); Wall(s) = Approx solver running time (seconds) on a modern CPU using the optimal parameters listed. Computational time to find optimal parameters is not taken into account; Vs. = Best algorithm for reverse engineering origami shape in terms of minimum *d*(**g, r**) achieved: (R)EVNANO, (N)AIVE or = (equal performance). Rows with ‘–’ entries signify that the origami could not be reverse engineered by the respective solver.

|  |  |  |  |  | NAIVE Solver |  |  |  | REVNANO Solver |  |  |  |  |  |  |  |
| --- | --- | --- | --- | --- | --- | --- | --- | --- | --- | --- | --- | --- | --- | --- | --- | --- |
| ID | Raster Origami | Ref | Dim | Staples | $\mu_{\min}$ | $d(\mathbf{g}, \mathbf{r})$ | SP% | Wall(s) | $\mu_{\min}$ | $\sigma$ | $\beta$ | $d(\mathbf{g}, \mathbf{r})$ | SP% | Wall(s) | Vs. | |
| 1 | RNA/DNA Hybrid Triangle | 24 | 2 | 22 | 6 | 16 | 100.0 | <1 | 6 | 0 | 0.10 | 16 | 100.0 | <1 | = |  |
| 2 | M1.3 Four Finger | 38 | 2 | 24 | 6 | 6 | 100.0 | <1 | 6 | 0 | 0.05 | 6 | 100.0 | <1 | = |  |
| 3 | Brick | 39 | 3 | 45 | 5 | 459 | 77.8 | <1 | 6 | 1 | 0.30 | 332 | 84.4 | <1 | R |  |
| 4 | Mini Triangle | 40 | 2 | 66 | 6 | 308 | 87.9 | <1 | 6 | 1 | 0.05 | 30 | 100.0 | 1 | R |  |
| 5 | DeBruijn Sequence Square | 41 | 2 | 70 | 6 | 44 | 100.0 | <1 | 6 | 0 | 0.05 | 44 | 100.0 | 1 | = |  |
| 6 | puc19 Rectangle | 33 | 2 | 90 | 6 | 2 | 100.0 | <1 | 6 | 0 | 0.30 | 2 | 100.0 | 1 | = |  |
| 7 | 6 Helix Bundle | 39 | 3 | 96 | – | – | – | – | 5 | 2 | 0.45 | 136 | 96.9 | 20 | R |  |
| 8 | Fivewell Plate | 42 | 2 | 158 | – | – | – | – | 6 | 6 | 0.30 | 100 | 100.0 | 272 | R |  |
| 9 | Membrane Nanopore | 43 | 3 | 172 | 6 | 2366 | 66.3 | 5 | 6 | 1 | 0.25 | 1275 | 83.7 | 140 | R |  |
| 10 | Single Staple Loop | 44 | 2 | 195 | 6 | 2 | 100.0 | 1 | 6 | 0 | 0.05 | 2 | 100.0 | 4 | = |  |
| 11 | DNA Frame | 45 | 2 | 222 | – | – | – | – | 6 | 3 | 0.55 | 174 | 99.5 | 71 | R |  |
| 12 | Rothmund Rectangle | 25 | 2 | 226 | – | – | – | – | 6 | 3 | 0.25 | 126 | 100.0 | 65 | R |  |
| 13 | Rectangle Variant | 46 | 2 | 226 | – | – | – | – | 6 | 3 | 0.30 | 114 | 100.0 | 57 | R |  |
| 14 | Small Moon | 47 | 2 | 234 | – | – | – | – | 5 | 2 | 0.05 | 258 | 99.1 | 367 | R |  |
| 15 | Rothmund Smiley | 25 | 2 | 243 | – | – | – | – | 5 | 2 | 0.15 | 172 | 99.2 | 199 | R |  |
| 16 | Rothmund Star | 25 | 2 | 244 | – | – | – | – | 6 | 4 | 0.45 | 116 | 99.6 | 110 | R |  |
| 17 | Capsule | 48 | 3 | 264 | – | – | – | – | – | – | – | – | – | – | – |  |
| ID | Wireframe Origami | Ref | Dim | Staples | $\mu_{\min}$ | $d(\mathbf{g}, \mathbf{r})$ | SP% | Wall(s) | $\mu_{\min}$ | $\sigma$ | $\beta$ | $d(\mathbf{g}, \mathbf{r})$ | SP% | Wall(s) | Vs. | |
| 18 | Triangle | 37 | 2 | 7 | 6 | 46 | 85.7 | <1 | 5 | 0 | 0.05 | 8 | 100.0 | <1 | R |  |
| 19 | Square | 37 | 2 | 8 | 6 | 6 | 100.0 | <1 | 6 | 0 | 0.05 | 6 | 100.0 | <1 | = |  |
| 20 | Pentagon | 37 | 2 | 10 | 6 | 6 | 100.0 | <1 | 6 | 0 | 0.05 | 6 | 100.0 | <1 | = |  |
| 21 | Tetrahedron | 37 | 3 | 13 | 5 | 141 | 76.9 | <1 | 5 | 0 | 0.05 | 4 | 100.0 | <1 | R |  |
| 22 | Triangle Mesh | 37 | 2 | 22 | 5 | 168 | 77.3 | <1 | 5 | 0 | 0.05 | 35 | 95.5 | <1 | R |  |
| 23 | Cube | 37 | 3 | 25 | 5 | 502 | 56.0 | <1 | 5 | 0 | 0.05 | 6 | 100.0 | <1 | R |  |
| 24 | Star Mesh | 37 | 2 | 38 | 6 | 443 | 76.3 | <1 | 5 | 0 | 0.05 | 66 | 97.4 | 1 | R |  |
| 25 | Dodecahedron | 37 | 3 | 59 | – | – | – | – | 5 | 0 | 0.10 | 188 | 96.6 | 3 | R |  |
| 26 | Icosahedron | 37 | 3 | 64 | – | – | – | – | 5 | 2 | 0.15 | 355 | 89.1 | 4 | R |  |
| 27 | Square Mesh 1 | 37 | 2 | 82 | – | – | – | – | 5 | 4 | 0.05 | 284 | 92.7 | 78 | R |  |
| 28 | Hexagon Mesh 1 | 37 | 2 | 84 | – | – | – | – | 5 | 2 | 0.25 | 335 | 92.9 | 13 | R |  |
| 29 | Annulus Mesh 1 | 37 | 2 | 89 | – | – | – | – | 5 | 0 | 0.05 | 91 | 98.9 | 7 | R |  |
| 30 | Hexagonal Tile | 37 | 2 | 98 | – | – | – | – | 5 | 0 | 0.05 | 169 | 98.0 | 12 | R |  |
| 31 | Truncated Cube | 37 | 3 | 98 | – | – | – | – | 6 | 2 | 0.05 | 72 | 100.0 | 6 | R |  |
| 32 | Cross Mesh | 37 | 2 | 110 | – | – | – | – | 5 | 1 | 0.15 | 176 | 97.3 | 18 | R |  |
| 33 | vHelix Ball | 49 | 3 | 132 | 6 | 115 | 98.5 | 1 | 6 | 0 | 0.20 | 18 | 100.0 | 5 | R |  |
| 34 | Annulus Mesh 2 | 37 | 2 | 161 | – | – | – | – | 5 | 3 | 0.05 | 106 | 100.0 | 131 | R |  |
| 35 | Lotus Mesh | 37 | 2 | 180 | – | – | – | – | 5 | 3 | 0.05 | 271 | 97.8 | 255 | R |  |
| 36 | Enneagonal Trapezohedron | 37 | 3 | 318 | – | – | – | – | – | – | – | – | – | – | – |  |

By contrast, Table I confirms that the NAIVE solver mentioned in the Introduction is not robust for recovering an origami contact map from raw sequences. A contact map could be reconstructed for only 44% of origamis tested, and in only 22% of origamis tested were ≥95% of staples placed. Moreover, for every origami, REVNANO consistently achieved an equal or lower base hamming distance *d*(**g, r**) to the ground truth contact map (final table column).

Note that in the above result statistics, ‘percent staples placed’ is a crude metric, as it simply reflects the total fraction of staples that were able to be routed by a solver. Some of these staples may have been routed incorrectly. Hamming distance is the most accurate metric to assess performance, but is less intuitive.

In general, performance of the REVNANO solver is sensitive to all three parameters: minimum staple leg size *μ*_min_, competitor routes allowed *σ*, and staple overlap tolerance *β*. Figure 3 exhaustively characterises performance across the smiley face parameter space for *μ*_min_ = 6bp. Parameter sensitivity typically varies significantly from origami to origami, with optimal performance wells located at different positions in the (*σ, β*) parameter space (see Supplementary Note 7 for examples). This is because local sequence configurations are determined by multiple factors, such as the particular origami scaffold and staple routing, as well as by the scaffold sequence itself and its rotation. In general, origamis displaying the highest parameter sensitivity are those with many short hybridising staple sections (e.g. 8bp or less) and/or a scaffold sequence with many repeated sequence regions. Conversely, origamis displaying low or no parameter sensitivity are those with long-leg staples (e.g. 16bp) and/or a scaffold with minimal repeats. Note that parameter *σ* typically has a sweet spot range: when too low, not enough competitor staples routes are considered and correct staple routes are omitted from staple routing trees; conversely, when *σ* is too high, staple routing trees contain a plethora of alternate staple routes and become either too large to construct, or constraint propagation at Stage 1 cannot narrow down staple routes effectively.

**FIG. 3.**
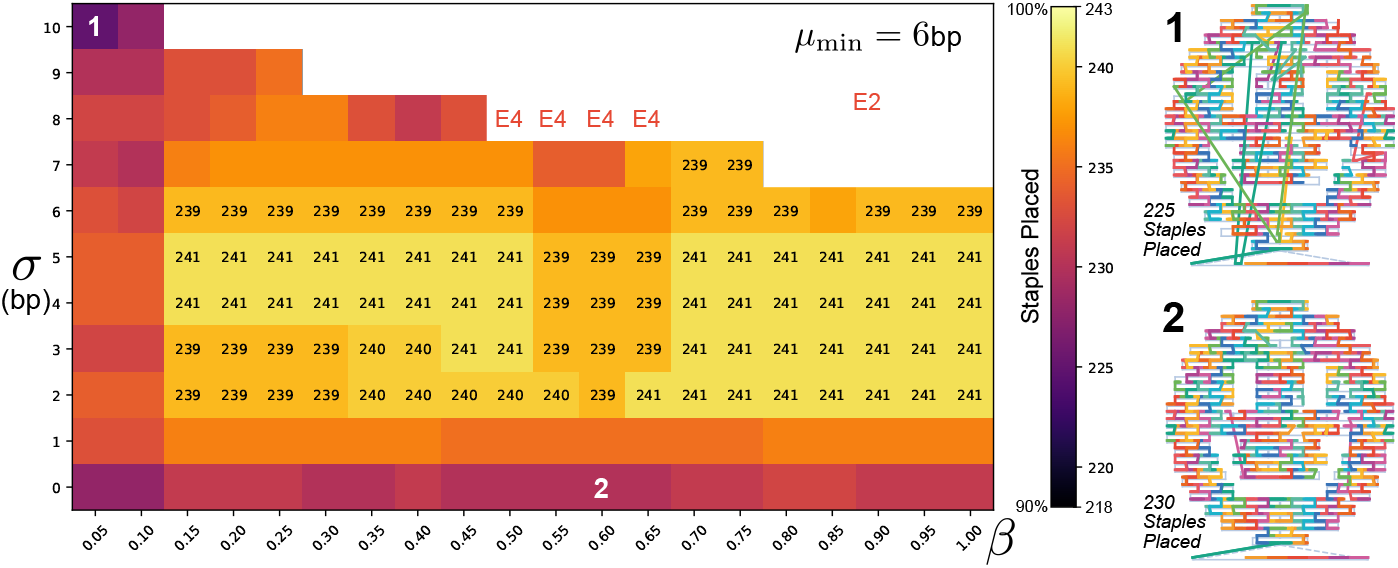
REVNANO Solver Parameter Sensitivity. Heatmap shows regions of the REVNANO parameter space (*μ*_min_ = 6bp, *σ, β*) where reverse engineering of the Rothemund Smiley^25^ contact map is most effective in terms of total number of staples placed. REVNANO running in deterministic staple placement mode. Smiley faces 1 and 2 (right) show how staple placement differs at (*σ* = 10bp, *β* = 0.05) and (*σ* = 0bp, *β* = 0.6) parameter points, respectively. White space at the top of the heatmap represents parameter points terminating in a REVNANO error condition: E2 = Not enough staples placed to perform reliable shortest-path calculations; E4 = Unresolvable 3-staple overlap exists. See Supplementary Note 7 for (*μ*_min_ = 5bp, *σ, β*) case, and REVNANO run times for each parameter combination.

While in the ideal case a grid search of parameter regimes would be conducted to find optimal REVNANO parameter combinations for a specific origami shape with specific sequences, in practice default ‘consensus’ parameters for raster and wireframe shapes can be derived (Supplementary Note 3). These default parameters yield acceptable REVNANO performance in most cases and are listed in Table II. Moreover, the consensus parameters represent a good starting point for parameter tweaking.

**TABLE II.** REVNANO Consensus Parameters.

| Origami Type | REVNANO Consensus Parameters |  |  |
| --- | --- | --- | --- |
| | $\mu_{\min}(\text{bp})$ | $\sigma(\text{bp})$ | $\beta$ |
| Raster | 6 | 4 | 0.25 to 0.3 |
| Wireframe | 5 | 2 | 0.2 to 0.25 |

It is instructive to now examine in detail some cases where REVNANO produces defective solutions, or fails to produce a solution at all, in order to better understand features and limitations of this solver.

### B. Cause of Minor Defects

The smiley face reverse engineered in Figure 2 ends up with five different minor defects (Figure 2f) which become evident when the recovered staple routings are super-imposed on the proper scaffold routing.

The defects are caused by the following reasons:

- In defect (1), the teal coloured staple is meant to crossover after 16 bases to pinch a loop of the M13 scaffold closed under the smiley face. However, this staple is actually complementary with the scaffold after the designed crossover point for 1 more base into the loop. Hence, sequence matching the staple (staple 5’ to 3’) with the scaffold causes the staple to be split 17-15 instead of 16-16. This is not picked up by Stage 3 overlap detection because the overshoot has not caused an overlap.
- In defects (2) and (3), an independent ‘C’ shaped edge staple should be hybridised at each of these locations. Instead a single staple incorrectly bridges both locations in the final REVNANO solution. This happens because of the following chain of events. The staple meant to be at (3), referred to as ‘staple (3)’ from now on, does not contain the correct designed route in its SRT because of sequence matching overshoot. The designed crossover point of this 12nt staple is 6-6, but the staple actually remains complementarity to the scaffold 2nt after the crossover point. Hence sequence matching splits staple (3) as 8-4 and the second section fails to be placed since 4 *< μ*_min_. However, staple (3) does have the bridge route in Figure 2f in its SRT. At iteration 1 in Figure 2d, when the orange staple is forced to its clearest shortest path, staple (3) happens to collapse to a single route, which is the latter incorrect bridge route, and it is placed. This staple placement precludes the staple meant to be at (2) from binding.
- In defect (4), a 3-section staple is missing because it fails to have a routing tree made at Stage 0. Meant to be split 8-16-8, this staple is split 9-18-5 by sequence matching overshoot and the final section of 5nt is too short to be matched (5 *< μ*_min_). Obviously, no other alternative routes exist on the scaffold for this particular staple, either.
- In defect (5), sequence matching overshoot (marked in red) causes the axis of the Holliday junction to be skewed toward the right by 1 base. Skewed Holliday junctions are in fact quite common place in the smiley face reconstruction and they increase the base hamming distance *d*(**g, r**) error.

Generally speaking, all defects of this smiley face example can be attributed to by-chance sequence matching overshoot at REVNANO Stage 0. In effect, this overshoot causes the constraint satisfaction problem to be improperly specified in the first instance, i.e. some staples do not contain a correct route in their SRT. The final defects are cascade effects from the original problem mis-specification. Sequence matching overshoot is, however, an artefact implicit in and non-separable from the sequence matching approach used by REVNANO. It can be concluded that a main challenge of the REVNANO solver is to first set up a correct description of the CSP problem which contains a final solution, rather than the process of actually solving the problem.

### C. Cause of Reverse Engineering Failure

Table I shows two instances in the test set (Origamis 17 and 36) for which a contact map cannot be reverse engineered under any REVNANO parameterisation.

The reason for REVNANO failing on Origami 17 (Capsule) is that this origami contains an unusually high number of staples (87) with very short (i.e. 3bp or less) hybridising sections. When staple routing trees are built at Stage 0, either these staples cannot be routed at all (since only sequence matches above *μ*_min_ are allowed in the SRT), or they are routed, but incorrectly, which in turn precludes other staples from assuming their correct route. The overall result is that, under any parameter setting, insufficient staples are placed in Stage 1 to allow reliable shortest path calculations to be performed at Stage 2 and the process fails. Many staples with short hybridising sections is in fact a sufficient condition for REVNANO failure, since the Capsule origami cannot have a contact map derived even when the scaffold sequence is changed for a repeat deficient order 7 DeBruijn sequence^41^. Of the origamis with less than 95% of staples placed by REVNANO (e.g. Origamis 3, 9, 26, 27, 28), most have a notable proportion of staples with extremely short hybridising sections.

REVNANO fails on Origami 36 (Enneagonal Trapezohe-dron) because this design features an unusually long 13530nt scaffold (nearly twice as long as most other designs tested) and the scaffold sequence additionally contains many repeated sequence regions. Staples on this origami have fairly long hybridising sections (the majority 10bp or more) and staple routing trees can be reliably constructed. However, the difficulty here is that each staple has a plethora of potential routes, due to both the scaffold length and number of repeated sequence regions. In parameter regime (*μ*_min_ = 6bp, *σ* = 5bp, *β* = 0.5) for example, 98% of staples have more than 1 route going into Stage 1, with an average number of 135 routes per staple. On the upper limit, one staple has 7829 possible scaffold routes. Such a high average number of routes per staple means that staple footprints are very small or non-existent, and thus constraint propagation is ineffective at Stage 1. Stage 2 subsequently raises the error that insufficient staples are placed to allow reliable shortest path calculations and REVNANO terminates. Interestingly, eliminating the scaffold repeats factor (by changing the scaffold sequence to an order 7 DeBruijn sequence with no repeated regions of 7nt or more) does reduce potential staple routes and allows recovery of a contact map.

Summarising the results so far, it can be rationalised that reduced REVNANO performance or failure is correlated with the following origami features: (i) Origamis with many staples containing very short (below *μ*_min_ threshold) hybridising sections causing staples to be routed incorrectly, or not routed at all; (ii) Scaffold sequences with many repeated regions, leading to false positive staple routes; (iii) Large origami designs (e.g. > 10000nt scaffold) where a longer scaffold increases the total number of staple routes possible; (iv) Origami designs with long ssDNA scaffold sections that permit uncon-tested yet incorrect staple routes; (v) The use of sub-optimal REVNANO parameter settings.

Typical REVNANO solve time ranged from a few seconds up to 6 minutes on a single modern CPU (with no optimisation of the underlying Python code). The main computational burden is the construction of the staple routing trees at Stage 0. When *μ*_min_ = 5bp, staple routing trees are wider than when *μ*_min_ = 6bp and Stage 0 time increases accordingly.

### D. Origami Guide Schematics

Figure 4 shows eight origamis reverse engineered through the full process of first recovering a contact map, and then creating a geometric guide schematic embedding this connectivity. Note that all guide schematics in Figure 4 are shown as static images, but they are actually dynamic HTML pages where origami shapes can be zoomed, rotated and re-arranged in 2D or 3D, and where mouse tooltip text provides part identification and sequence information while moving over the schematic.

**FIG. 4.**
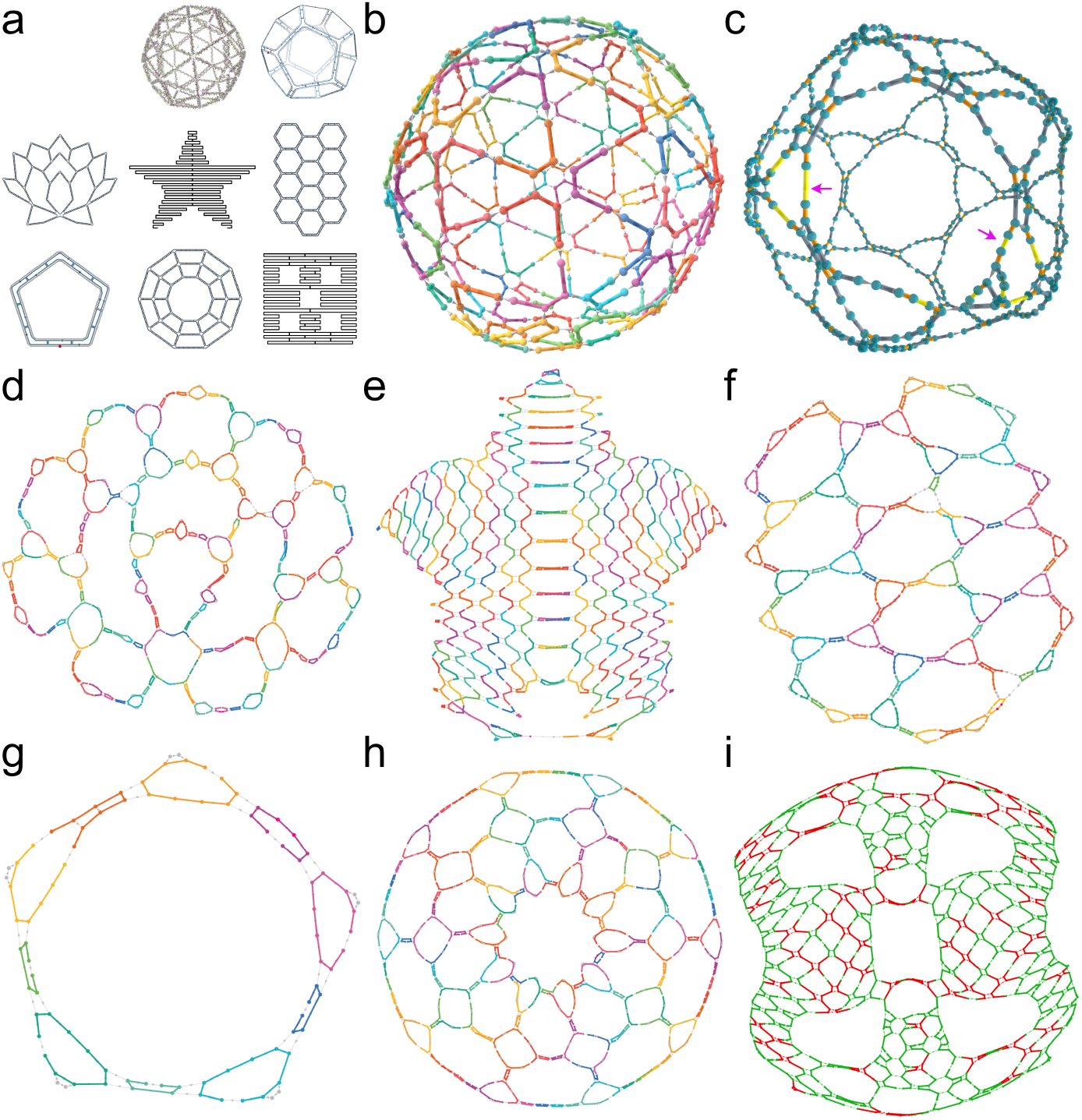
Example Origami Guide Schematics Reverse Engineered from Raw Scaffold and Staple Sequence Lists. **(a)** Original origami schematics. Raw sequences derived from these schematics were passed through the reverse engineering pipeline (Figure 1b) using REVNANO with optimal parameter settings, to produce the following reconstructed guide schematics: **(b)** 3D Ball in staples view, 100% staples placed; **(c)** 3D Dodecahedron in scaffold routing view, 96.6% staples placed (purple arrows highlight regions where staples omitted); **(d)** 2D Lotus Mesh in staples view, 97.8% staples placed; **(e)** 2D Rothemund Star in staples view, 99.6% staples placed; **(f)** 2D Hexagonal Tile in staples view, 98% staples placed; **(g)** 2D Pentagon in staples view, 100% staples placed; **(h)** 2D Annulus Mesh 2 in staples view, 100% staples placed; **(i)** 2D Fivewell Plate, 100% staples placed shown in sequence-ambiguous junctions view computed by the AMBIG algorithm: immoveable junctions in green, moveable junctions in red. When using the Fivewell Plate guide schematic to re-enter the design into an origami CAD tool, symmetry can be used to guess the correct positions of the ambiguous red crossovers by looking at the positions of the unambiguously positioned green crossovers. Guide schematics for all other origamis in the test set of Table 1 are given in Supplementary Note 8.

We found that guide schematics were generally intelligible for all reverse engineered 2D origamis and for small 3D wireframe origamis which formed compact shapes like cubes and spheres (Figure 4b and c).

Conversely, we observed that guide schematics for large raster 3D shapes, like the Brick, 6-Helix Bundle, and Membrane Nanopore (Origamis 3, 7, and 9 respectively) were not intelligible. Raster 3D shapes have many dense internal connections which make it hard to discern staple routings, even with an interactive schematic. Also, the layout task for the spring embedder is less constrained in 3-dimensions with there being 2 extra degrees of freedom in comparison to 2-dimensions. This extra freedom tends to destroy the form of 3D shapes which are not tightly-constrained by their geometry. For the same reason, we found that omitted staples have a stronger negative effect on reconstructed 3D schematics.

Generally, we found the spring embedder algorithm to have a graceful degradation in geometric layout as staples were omitted from an origami design. However, in two cases in 2-dimensions (Square Mesh 1 and Hex Mesh 1) the spring embedder suddenly shifted from a good layout at 100% staples placed to a poor (locally optimal) crossed layout at 93% staples placed.

Overall, of 34 guide schematics reconstructed from the origami test set, 29 could be judged of sufficient quality to assist the manual re-input of the origami design into an origami CAD tool (see Supplementary Note 8).

One striking feature of geometry re-constructed by spring-layout is that the rectilinear cells of an origami schematic have an inflated ‘chicken wire’ appearance. This is a result of the Kamada-Kawai spring layout algorithm trying to make the Euclidean distance between nodes equal to the shortest path (Manhattan) distance. A similar inflation effect is observed at the multi-arm junctions of the ATHENA wireframe shapes. There likely exist solutions to restore more rectilinearity to the geometry (see Discussion), but the organic spring-layout does have the advantage that holes naturally appear in shapes where only helix stacking interactions and not base pair hydrogen bonding hold the structure in place. From a design perspective, these areas can be useful to identify.

### E. Effect of Input Sequence Noise

Finally, when noise was added to input staple sequences, as could be the case when sequences are imperfectly copied from a publication, it was found that REVNANO performance generally decreased (see Supplementary Note 5 for detailed analysis). Interestingly however, the solver could still often reconstruct the correct origami contact map when the staple sequence pool was ‘contaminated’ by surplus random DNA staples, particularly when the surplus sequences were added at the end of the correct staple list.

## IV. DISCUSSION

In this work, we developed a two-stage computational pipeline for reverse engineering an origami nanostructure guide schematic from a list of origami staple sequences (plus the scaffold sequence). In the first stage of our pipeline, we demonstrated that the REVNANO solver was robust at recovering a contact map for origamis even with challenging features such as repeated scaffold sequence regions and staples with short (e.g. 7 or 8bp) binding sections. In the second stage of our pipeline, we also demonstrated that spring-graph layout is an effective procedure for reconstructing origami geometry from topology/connectivity. Overall, from the 36 origamis drawn from the literature, we were able to obtain an intelligible guide schematic for 29 of them, which included all 2D origami shapes and those 3D shapes which were compact wireframe designs. Therefore, overall, we succeeded at converting unstructured origami sequence data back into a meaningful geometric form.

### A. Limitations

The main limitations of our approach, with some ideas for mitigation are as follows. Firstly, the solution accuracy of REVNANO is critically dependent on the (*μ*_min_, *σ, β*) parameters of the solver. It is sometimes necessary to conduct a scan of parameter space in order to arrive at optimal performance. However, we did derive a set of consensus parameters for both raster and wireframe origami shapes which can serve as good starting points in this parameter search.

3D raster origamis are typically hard to reverse engineer as the guide schematic ends up with dense internal connectivity which is hard to decipher. Also, some 3D shapes are poorly constrained in their reconstructed spring-layout geometry and feature e.g. kinks which are not in the original origami design, for example with the 6 Helix Bundle (Origami 7) shown in Supplementary Note 8. This is an inherent limitation with the spring-layout reconstruction used.

Another limitation is that misplaced staples are sometimes an undesired output of the solver (e.g. in the Smiley example, Figure 2d). Without knowing the *a priori* scaffold routing of an origami nanostructure, misplaced staples are hard to detect visually as they just manifest as complicated tangled regions of the final origami guide schematic. One strategy to detect misplaced staples may be to run the REVNANO solver twice, in two opposite sequence matching directions, one matching staples 5’ to 3’ (as is implemented) and another matching staples 3’ to 5’ instead. Only those staples placed consistently in each matching direction would be included in the final guide schematic. Another strategy to identify/eliminate misplaced staples could be to calculate the distribution of staple crossover lengths for all placed staples and eliminate those staples with large outlier values. This distribution would be constructed by considering each staple, removing each crossover in turn, and calculating the shortest path through the origami mesh connecting the sides of the respective staple crossover in its absence. However, a risk with this procedure is that false positives like staples closing a flat sheet into a tube could also be eliminated.

A final limitation is that non-hybridising regions of staples like dangles or interior loopout regions need pre-marking in the sequence input file to REVNANO. As discussed in more detail in Supplementary Note 6, the solver cannot automatically detect these regions. Therefore, some meta-markup in the raw staple sequences is required for some origami shapes (notably wireframe origami shapes). Although non-hybridising staple regions tend to be polyT, detecting these regions is not trivial since polyT regions can also legitimately exist in hybridised sections of staples.

### B. Future Directions

In the short term, one improvement to our approach could be to force the origami guide schematic to assume a more rectilinear geometry by diagonally bracing cells with invisible links in the origami graph 𝔻. This approach would make Manhattan distances between two points on the nanostructure even more closely approximate the actual Euclidean distance and reduce the inflated appearance of individual cells. The challenge, of course, is to develop an algorithm able to robustly identify cells in an origami schematic when features like complex multi-arm junctions, or even circular geometry^50^, may be present.

The most obvious extension to our reverse engineering pipeline would be to automatically derive an editable caDNAno (or scadnano) schematic from the guide schematic of the origami. It could be envisioned that e.g. a branch and bound search could tie the origami graph D to a compatible square/honeycomb caDNAno lattice representation, perhaps using orientations of staple crossovers etc. on the geometric guide schematic as search heuristics. However, while an interesting possibility, this transformation from guide schematic to (s)caDNAno turns out to be possible only for a very small subset of origami nanostructures. The transformation essentially requires that information loss *I*_1_ in Figure 1a is close to zero, which requires that an origami design meet a very strict set of conditions, such as: (i) the existence of a single large connected component on the design grid with no long-range crossovers connecting non-adjacent parts of the design (ii) no base insertions or base deletions; (iii) no single stranded scaffold regions, for example at the end of helix rows or in the form of a large loop under the shape; (iv) all junctions rigidly fixed in place by tight sequence constraints, and so on. In the general case, therefore, the best that can be reliably achieved is the automatic creation of an (off-grid) origami guide schematic which is then manually turned into an origami CAD schematic via the general intelligence and stylistic choices of a human.

The most promising future extension to our work may be to directly convert the geometric layout of origami graph D into the oxDNA format^31^, allowing the guide schematic to be opened in (grid free) tools such as oxView^32^ for base editing and for direct coarse-grain dynamics simulation. This would involve the replacement of ssDNA and dsDNA graph edges with random single stranded DNA regions and standard double helices, respectively, to give an initial non-relaxed oxDNA configuration. In the best case, this feature would allow an origami structure specified only as sequences (and lacking a schematic) to be relaxed and have its equilibrium properties computed nevertheless.

### C. General Comments

To close with some general comments, it is worth mentioning that a topological origami contact map - without a geometric representation - can be useful in its own right. For example, it can be used to generate new complementary staple sequences for an origami given a change of scaffold sequence. In future data science applications, origami contact maps could be compared according to some distance measure to infer which origamis may have similar folding characteristics. Alternatively the domain-level graph of the origami (equivalent to the contact map) could be used to provide origami connectivity information which could provide extra input features for future machine learning models predicting e.g. optimal self-assembly protocols. The domain-level graph is also useful for coarse-grain kinetic simulations of origami self-assembly^33^ or dis-assembly. Usefully, the contact map for an origami can be unambiguously encoded into the staple and scaffold sequences of a nanostructure (i.e. *I*_2_ = 0 loss in Figure 1a) if (i) REVNANO places all staples and (ii) no junctions have sequence ambiguity. The latter property can be trivially engineered by ensuring that the scaffold base letter following every staple crossover point is always different to the scaffold base letter on the other side of the staple crossover.

Finally, we expect a crude correlation between those origami nanostructures which reverse engineer quickly and completely, and those that self-assemble with few potential kinetic traps. This is because ease of reverse engineering is associated with staples that have unique, or close to unique, scaffold routes. The correlation can only be expected to be crude, however, because (i) a REVNANO staple routing tree contains only a subset of all possible binding possibilities for a staple (partial staple routes are omitted) and (ii) strand displacement and scaffold secondary structure are absent in REVNANO solution construction but are important processes operating in physical origami self-assembly.

## V. CONCLUSION

In this study, we proposed a first algorithmic solution to the inverse problem of converting raw staple and scaffold sequence lists back to a guide schematic for an origami nanostructure. We demonstrated good performance of our pipeline on a test set of 36 origamis from the literature.

Reverse engineering is not an exact science and success in rebuilding original source data critically depends on how much information is lost in the conversion process to final artefact. In the case of origami nanostructure design, we were able to partially recover two main stages of information loss to an adequate extent as to rebuild an approximate origami guide schematic in most cases. By integrating further information such as ambiguity of junctions, symmetries in the global origami design, and other considerations, human general intelligence is then able to make the final bridge back to an editable source schematic in an origami CAD design tool of choice.

In general, our work helps to make the exchange of origami nanostructure designs in nanobiotechnology more transparent and reliable, if raw sequence lists continue to be the de facto exchange format. Also, through the creation of contact maps from origami sequences mined from publications, our work can contribute to future data-driven approaches in the nucleic acid nanotechnology field.

## CODE AVAILABILITY

Python source code (MIT licence) and documentation is located at https://revnano.readthedocs.io. The code includes a Python JupyterLab notebook for reverse engineering origami schematics from sequence lists.

