## Supplementary Information for "Reverse Engineering DNA Origami Nanostructure Designs from Raw Scaffold and Staple Sequence Lists"

### Contents

**Supplementary Note 1**
**Origami Contact Map and Origami Domain-Level Graph** **2**

**Supplementary Note 2**
**Derivation of Ground Truth Origami Contact Maps** **6**

**Supplementary Note 3**
**REVNANO Consensus Parameters** **9**

**Supplementary Note 4**
**REVNANO in Non-Deterministic Mode** **10**

**Supplementary Note 5**
**REVNANO with Noisy Staple Sequences** **13**

**Supplementary Note 6**
**REVNANO with Staple Dangles and Staple Loopouts** **20**

**Supplementary Note 7**
**REVNANO Parameter Space Examples** **22**

**Supplementary Note 8**
**Reverse Engineered Guide Schematics** **25**

### Supplementary Note 1 Origami Domain-Level Graph and Origami Contact Map

Supplementary Figure 1 makes explicit the relation between an example origami schematic drawn in scadnano[1], the origami domain-level graph $\mathbb{D}$ and the origami contact map. Even though REVNANO creates the contact map first and the domain-level graph later, it is most instructive to discuss them in the opposite order.

In the domain-level graph $\mathbb{D}$, nodes correspond to individual bases on the scaffold strand, and vertices represent how these bases are connected, either through hybridised double stranded DNA helices (scaffold hybridised to a staple leg), single stranded scaffold DNA, scaffold nicks (**n**), staple crossovers (**x**), staple loopouts (**lo**) or staple dangling ends (**d**).

Supplementary Figure 1c shows a domain-level graph $\mathbb{D}$ for a simple example origami, drawn in two different representations for convenience. Panel c(i) shows the scaffold strand routing view with scaffold domains numbered from 0 starting at scaffold 5'. Panel c(ii) highlights staple routings instead, with blue numbers on graph nodes indicating the scaffold base number, again numbered from 0 starting at scaffold 5'. The domain-level graph contains all information from both representations. Note that both single-stranded regions and double-stranded helices are always represented by *single* vertices in the graph. Supplementary Figure 2 outlines the solution to a technical side case sometimes arising at wireframe origami junctions where a staple makes a bulge loop with the scaffold. In this case, two vertices are simultaneously possible between adjacent nodes in the domain-level graph.

Any origami nanostructure can be converted into a topological domain-level graph representation, with the exception of origamis which have scaffold domains of a single base (see Supplementary Figure 1b). The minimum length of a ssDNA or dsDNA domain on the domain-level graph is 2nt, since every domain vertex must connect two nodes with different scaffold base indexes. Note that scaffold crossovers (i.e. where a significant re-orientation of the scaffold strand takes place) are not a distinct vertex type on the domain-level graph. In general, scaffold crossovers only become obvious in the context of the whole geometric layout of an origami. Thus, on the topological domain-level graph, scaffold crossovers are not differentiated from scaffold nicks (**n**).

The contact map representation of the origami (Supplementary Figure 1d) specifies which staple and scaffold bases are hybridised in the origami design. The contact map is split into two parts, both of which are necessary. The first part (Supplementary Figure 1d(i)) specifies how staples are hybridised with the scaffold:

- `staple_id` is the number of the staple
- `sequence` is the staple sequence, going from staple 5' to staple 3'
- `num_sections` is the number of distinct sequence regions on the staple: some staple sections will hybridise to domains on the scaffold, other staple sections will be single stranded dangles or single stranded interior loopouts not hybridised with the scaffold
- `section_lengths` are the number of nucleotides in each staple section (semicolon separated list)
- `section_dom_id` is the scaffold domain number that each staple section hybridises to (semicolon separated list). Scaffold domain numbers are shown on Supplementary Figure 1c(i). If a staple section does not hybridise to the scaffold, -1 is listed

- scaffold_base_ids are the scaffold base indexes that each base of the staple hybridises to, going from staple 5’ to staple 3’ (semicolon separated list). Scaffold base indexes are shown on Supplementary Figure 1c(ii). If a staple base does not hybridise with the scaffold, -1 is listed

The second part of the contact map (Supplementary Figure 1d(ii)) specifies how the scaffold is hybridised to staples:

- scaffold_base_id is the index of the scaffold base, beginning 0 at scaffold 5’ and increasing toward scaffold 3’
- base is the nucleotide base at scaffold_base_id
- dom_id is scaffold domain that this scaffold base is part of (numbers on Supplementary Figure 1c(i))
- staple_id is the staple that this scaffold base is hybridised to. If the scaffold base is not hybridised to a staple, -1 is listed
- staple_section_id is the staple section on staple staple_id that this scaffold base is hybridised to (first section at staple 5’ is numbered 0). If the scaffold base is not hybridised to a staple, -1 is listed
- staple_base_id is the staple base number on staple staple_id that this scaffold base is hybridised to (first staple base at staple 5’ is numbered 0). If the scaffold base is not hybridised to a staple, -1 is listed

Note that the first part of the contact map (staples to scaffold mapping) only completely specifies the second part of the contact map (scaffold to staples mapping) when all scaffold bases are hybridised with a staple, i.e. when the scaffold has no single stranded regions like surplus scaffold loops, or polyT regions at origami shape edges to prevent origami-origami stacking interactions. Similarly, the second part of the contact map only completely specifies the first part when all staple bases are hybridised with the scaffold, i.e. when staples have no single stranded dangle sections or loopout sections. In the general case, both parts of the contact map are required, neither is redundant.

Any origami nanostructure can be converted to a contact map, even origamis with 1nt scaffold domains (Supplementary Figure 1b). To make a domain-level graph from a contact map with 1nt (unhybridised) domains present, either 1nt domains can be removed by deleting the offending scaffold base, or surrounding staples can be removed. 1nt unhybridised domains can often occur at junctions in wireframe origami shapes, or at sites where REVNANO has made a subtle staple placement error.

A contact map can also represent the pathological case of a hybridised single nucleotide domain on the scaffold.

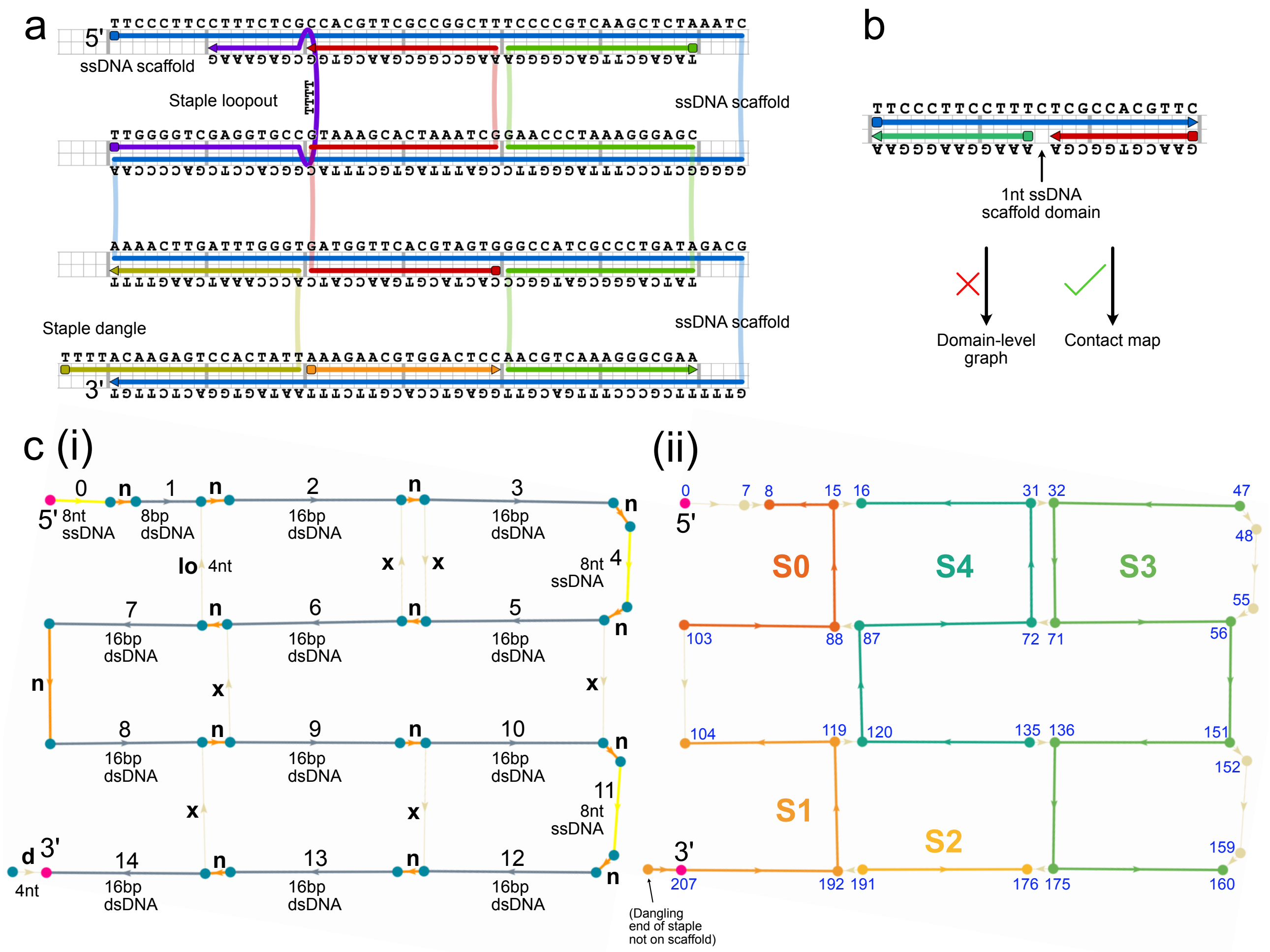


d

#### (i) Staples ⟶ Scaffold Mapping

staple_id ↓

| | sequence | num_sections | section_lengths | section_dom_id | scaffold_base_ids |
|---|---|---|---|---|---|
| 0 | TTGGGGTCGAGGTGCCTTTTCGAGAAAG | 3 | 16;4;8 | 7;-1;1 | 103;102;101;100;99;98;97;96;95;94;93;92;91;90;89;88;-1;-1;-1;-1;15;14;13;12;11;10;9;8 |
| 1 | TTTTACAAGAGTCCACTATTACCCAAATCAAGTTTT | 3 | 4;16;16 | -1;14;8 | -1;-1;-1;-1;207;206;205;204;203;202;201;200;199;198;197;196;195;194;193;192;119;118;117;116;115;114;113;112;111;110;109;108;107;106;105;104 |
| 2 | AAAGAACGTGGACTCC | 1 | 16 | 13 | 191;190;189;188;187;186;185;184;183;182;181;180;179;178;177;176 |
| 3 | TAGAGCTTGACGGGGAGAACCCTAAAGGGAGCTATCAGGGCGATGGCCAACGTCAAAGGGCGAA | 4 | 16;16;16;16 | 3;5;10;12 | 47;46;45;44;43;42;41;40;39;38;37;36;35;34;33;32;71;70;69;68;67;66;65;64;63;62;61;60;59;58;57;56;151;150;149;148;147;146;145;144;143;142;141;140;139;13 |
| 4 | CACTACGTGAACCATCGTAAAGCACTAAATCGAAGCCGGCGAACGTGG | 3 | 16;16;16 | 9;6;2 | 135;134;133;132;131;130;129;128;127;126;125;124;123;122;121;120;87;86;85;84;83;82;81;80;79;78;77;76;75;74;73;72;31;30;29;28;27;26;25;24;23;22;21;20;1 |

#### (ii) Scaffold ⟶ Staples Mapping

scaffold_base_id ↓

| | base | dom_id | staple_id | staple_section_id | staple_base_id |
|---|---|---|---|---|---|
| 0 | T | 0 | -1 | -1 | -1 |
| 1 | T | 0 | -1 | -1 | -1 |
| 2 | C | 0 | -1 | -1 | -1 |
| 3 | C | 0 | -1 | -1 | -1 |
| 4 | C | 0 | -1 | -1 | -1 |
| 5 | T | 0 | -1 | -1 | -1 |
| 6 | T | 0 | -1 | -1 | -1 |
| 7 | C | 0 | -1 | -1 | -1 |
| 8 | C | 1 | 0 | 2 | 27 |
| 9 | T | 1 | 0 | 2 | 26 |
| 10 | T | 1 | 0 | 2 | 25 |
| 11 | T | 1 | 0 | 2 | 24 |
| 12 | C | 1 | 0 | 2 | 23 |
| 13 | T | 1 | 0 | 2 | 22 |
| 14 | C | 1 | 0 | 2 | 21 |
| 15 | G | 1 | 0 | 2 | 20 |
| 16 | C | 2 | 4 | 2 | 47 |
| 17 | C | 2 | 4 | 2 | 46 |
| 18 | A | 2 | 4 | 2 | 45 |
| 19 | C | 2 | 4 | 2 | 44 |
| 20 | G | 2 | 4 | 2 | 43 |
| 21 | T | 2 | 4 | 2 | 42 |
| 22 | T | 2 | 4 | 2 | 41 |
| 23 | C | 2 | 4 | 2 | 40 |
| 24 | G | 2 | 4 | 2 | 39 |
| 25 | C | 2 | 4 | 2 | 38 |

Supplementary Figure 1: (a) Origami schematic ⟺ (c) origami domain-level graph $\mathbb{D}$ ⟺ (d) origami contact map relationship. Example origami for illustration purposes only. Panel (b) shows a 1nt scaffold domain which cannot be represented in a domain-level graph, but can be represented in a contact map. See text.

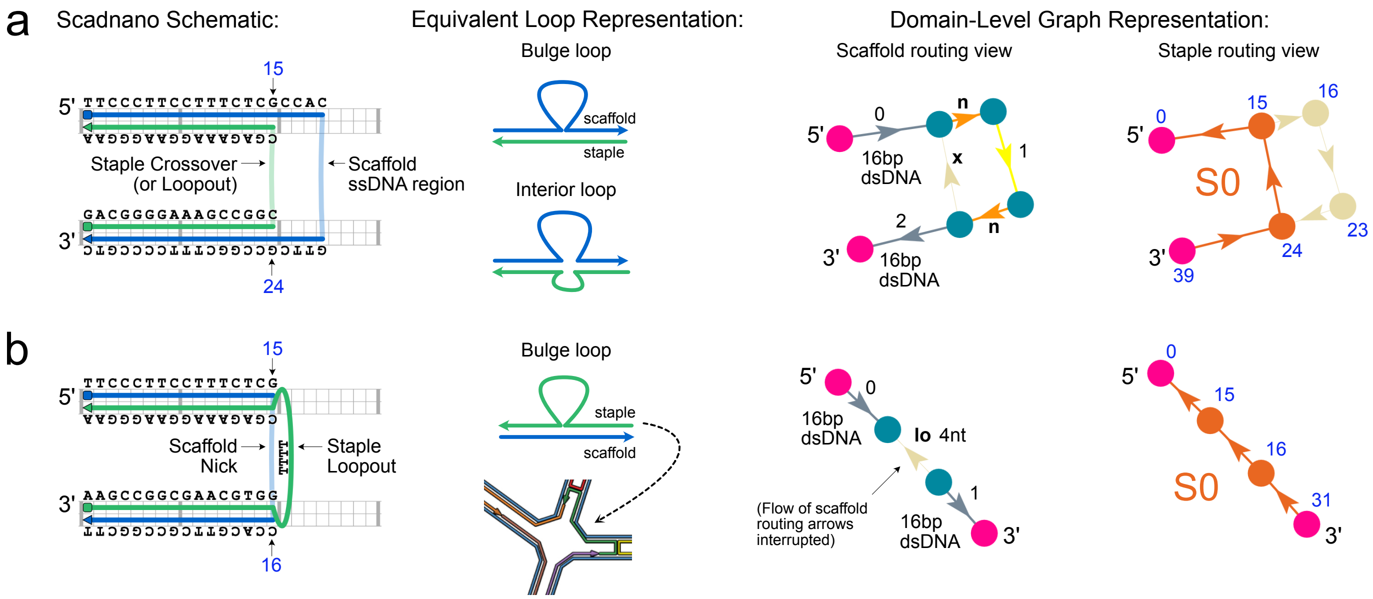


Supplementary Figure 2: When competing vertices arise in a domain-level graph. (a) No competing vertices case. A scaffold ssDNA region forms a bulge loop with a staple, or forms an interior loop with a staple loopout. Both the scaffold ssDNA region and staple crossover can be represented as separate vertices on the origami domain-level graph. (b) Competing vertices case. A staple loopout forms a bulge loop with the scaffold. This case often exists inside multi-arm junctions of wireframe origamis. Either the scaffold nick or the staple loopout could be an edge in the domain-level graph, linking scaffold bases 15 and 16. In this case, we use the edge for the staple loopout in the domain-level graph, giving it priority over the edge for the scaffold crossover. A side effect is that the staple loopout vertex interrupts the flow of the scaffold routing arrows in the origami guide schematic.

### Supplementary Note 2 Derivation of Ground Truth Origami Contact Maps

| ID | Origami | Ground Truth Contact Map Obtained via Route |
|---|---|---|
| 1 | RNA/DNA Hybrid Triangle | 1 |
| 2 | M1.3 Four Finger | 1 |
| 3 | Brick | 2 |
| 4 | Mini Triangle | 2 |
| 5 | DeBruijn Sequence Square | 2 |
| 6 | puc19 Rectangle | 1 |
| 7 | 6 Helix Bundle | 2 |
| 8 | Fivewell Plate | 1 |
| 9 | Membrane Nanopore | 2 |
| 10 | Single Staple Loop | 1 |
| 11 | DNA Frame | 1 |
| 12 | Rothemund Rectangle | 1 |
| 13 | Rectangle Variant | 1 |
| 14 | Small Moon | 2 |
| 15 | Rothemund Smiley | 1 |
| 16 | Rothemund Star | 1 |
| 17 | Capsule | 2 |
| 18 | Triangle | 3 |
| 19 | Square | 3 |
| 20 | Pentagon | 3 |
| 21 | Tetrahedron | 3 |
| 22 | Triangle Mesh | 3 |
| 23 | Cube | 3 |
| 24 | Star Mesh | 3 |
| 25 | Dodecahedron | 3 |
| 26 | Icosahedron | 3 |
| 27 | Square Mesh 1 | 3 |
| 28 | Hexagon Mesh 1 | 3 |
| 29 | Annulus Mesh 1 | 3 |
| 30 | Hexagonal Tile | 3 |
| 31 | Truncated Cube | 3 |
| 32 | Cross Mesh | 3 |
| 33 | vHelix Ball | 4 |
| 34 | Annulus Mesh 2 | 3 |
| 35 | Lotus Mesh | 3 |
| 36 | Enneagonal Trapezohedron | 3 |

Supplementary Table 1: Route used to derive a ground truth contact map for each origami (see following page).

Note: scripts `scadnano2contact.py`, `oxdna2contact.py` and `contactutils.py` referenced below are part of the origami contact map Python module released with this paper, available at:
https://contactmap.readthedocs.io

**Route 1**

1. The origami schematic was re-drawn by hand in scadnano[1] on a square grid, using schematic images in the original publication supplementary material. Any loop under the shape was also included

2. The original published scaffold sequence was applied to the design, and the `.sc` file exported (containing sequences)

3. Python script `scadnano2contact.py` converted the `.sc` file to a contact map (CSV format)

4. The scaffold was joined into a circle by Python script `contactutils.py`, if necessary

**Route 2**

1. A caDNAno[2] origami schematic (`.json`) was obtained from the original author, or from a publicly available repository

2. TacoxDNA[3] was used to convert caDNAno `.json` format to oxDNA format (topology and configuration files), using the appropriate square or hexagonal grid. TacoxDNA assigned a random scaffold sequence and complementary staple sequences

3. Python script `oxdna2contact.py` converted the oxDNA files to a contact map (CSV format)

4. The published scaffold sequence was applied to the contact map using Python script `contactutils.py`. This changed all staple sequences to be complementary to the new scaffold sequence

5. The scaffold was joined into a circle by Python script `contactutils.py`, if necessary

**Route 3**

1. Origami was exported from the ATHENA UI tool[4]. Export included a caDNAno `.json` file on hexagonal grid with no sequence information included. All scaffolds were linear (non-circular)

2. TacoxDNA[3] was used to convert caDNAno `.json` format to oxDNA format (topology and configuration files). TacoxDNA assigned a random scaffold sequence and complementary staple sequences

3. Python script `oxdna2contact.py` converted the oxDNA files to a contact map (CSV format)

4. The scaffold sequence listed in the `TXT_Sequence.txt` file output by ATHENA was applied to the contact map, using Python script `contactutils.py`. This changed all staple sequences to be complementary to the new scaffold sequence

5. Positions of staple loop-outs and dangles in the contact map were verified as being in identical positions as * delimited regions in staple sequences in the `TXT_Sequence.txt` file output by ATHENA

**Route 4**

1. Autodesk Maya (vHelix) source file was obtained from `http://www.vhelix.org/`
2. TacoxDNA[3] was used to convert vHelix format to oxDNA format (topology and configuration files). TacoxDNA assigned a random scaffold sequence and complementary staple sequences
3. Python script `oxdna2contact.py` converted the oxDNA files to a contact map (CSV format)
4. The published scaffold sequence was applied to the contact map using Python script `contactutils.py`. This changed all staple sequences to be complementary to the new scaffold sequence
5. The scaffold was joined into a circle by Python script `contactutils.py`

### Supplementary Note 3 REVNANO Consensus Parameters

The procedure below was used to derive REVNANO consensus parameters for raster and wireframe origami classes.

A base hamming distance heatmap was created for each origami (like for the origamis in Supplementary Figures 9 and 10). Base hamming distances were normalised by dividing by the origami scaffold length, giving a heatmap showing the percent fraction of incorrectly assigned scaffold bases. Normalisation was necessary such that larger origamis with larger absolute base hamming distances did not dominate cumulative scores. All percent fraction heatmaps for raster origamis with 95% or more staples placed were super-imposed and added by matrix addition. Similarly, all percent fraction heatmaps for wireframe origamis with 95% or more staples placed were super-imposed and added by matrix addition. This was done for $\mu_{\min} = 5$bp and $\mu_{\min} = 6$bp for both origami classes to obtain the results in Table 2 of the paper.

Optimal REVNANO parameters for an origami shape depend on a variety of factors like scaffold and staple routing, and the scaffold sequence used. The majority of raster origamis in Table 1 of the paper have a scaffold sequence based on M13. Likewise, the ATHENA tool exports wireframe origamis with a M13mp18 scaffold up to 7249nt and a Lambda phage sequence for larger designs[4]. Therefore, as scaffold sequences have approximately the same sub-sequence repeat properties in both classes of origamis, the qualitative difference in optimal parameters may stem from differences in staple routing: a staple in a typical raster origami has more crossovers to different helices and also more immediate staple neighbours than a staple in a typical wireframe origami (Supplementary Figure 3).

Note that the consensus parameters are also expected to perform well when scaffold sequences are synthetic with less repeats (i.e. DeBruijn sequence). In this case, REVNANO only has weak dependence on parameters and most parameter combinations give good results.

Supplementary Table 2 shows REVNANO performance on all origamis, using consensus parameters.

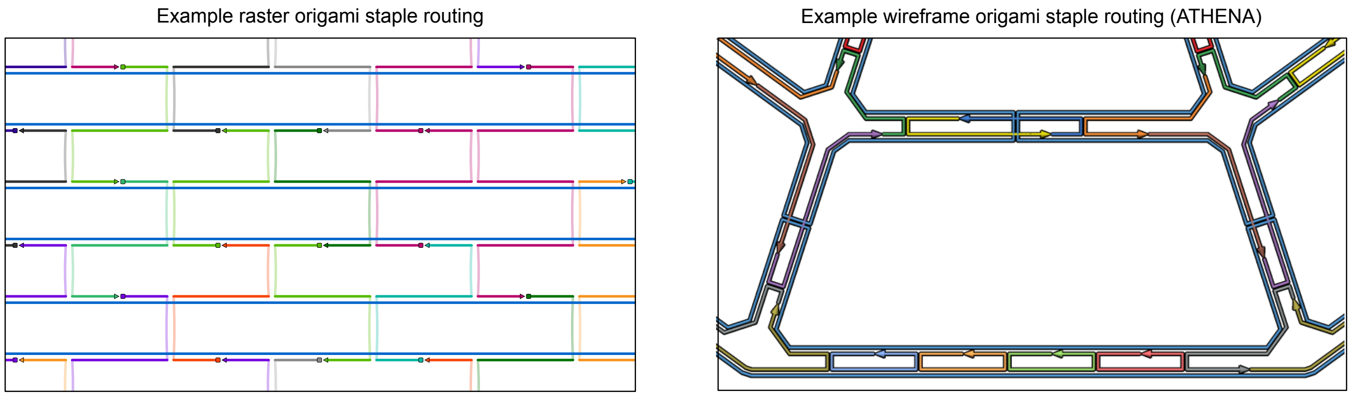


Supplementary Figure 3: Qualitative differences in staple routing between raster and wireframe origamis.

### Supplementary Note 4 REVNANO in Non-Deterministic Mode

REVNANO can be run in two modes, deterministic or non-deterministic:

- In deterministic mode the REVNANO solver polls staples (at Stages 0, 1 and 2) in the same order they are listed in the input sequence file
- In non-deterministic mode the solver polls staples in a random order

It may be hypothesised that the REVNANO solver could reconstruct different origami contact maps under different staple poll orders, because staple poll order changes the order in which constraints become established in the solving process. That is, if some staples are placed in certain ways first, they may prohibit other staples being placed in certain ways later.

However, Supplementary Table 2 shows that, in practice, this reasoning is only partially true. Yes, staple poll order can make a difference to the REVNANO solver in some cases: for 9 of 33 origamis tested, putting REVNANO in non-deterministic mode and running many times ($n = 250$ samples) can reach better solutions[1] than deterministic mode. However, as Supplementary Table 3 highlights, these solutions are only marginally better. Hamming distance only decreases by a few tens of bases, and staples placed only increases by around 1%, with respect to deterministic mode. For the majority of origamis tested, many samples in non-deterministic mode create exactly the same contact map as a single run in deterministic mode does (i.e. staple poll order makes no difference to the final contact map reconstructed).

In conclusion, rather than sampling many different staple poll orders in non-deterministic mode, compute effort seems better spent using the deterministic staple poll order and instead searching for REVNANO parameter combinations $(\mu_{\min}, \sigma, \beta)$ which give optimal performance. This is why REVNANO is used in deterministic mode in the paper.

[1] Where "better solution" means lower hamming distance to ground truth contact map, or more staples placed, or both.

| | | | | | Deterministic | | Non-deterministic | | | | | | |
|---|---|---|---|---|---|---|---|---|---|---|---|---|---|
| | | | | | $d(\mathbf{g},\mathbf{r})$ | SP% | | $d(\mathbf{g},\mathbf{r})$ | | | SP% | | |
| ID | Origami | $\mu_{\text{min}}$ | $\sigma$ | $\beta$ | | | $n$ | min | mean | max | min | mean | max |
| 1 | RNA/DNA Hybrid Triangle | 6 | 4 | 0.30 | 16 | 100.00 | 250 | 16 | 16.00 | 16 | 100.00 | 100.00 | 100.00 |
| 2 | M1.3 Four Finger | 6 | 4 | 0.30 | 6 | 100.00 | 250 | 6 | 6.00 | 6 | 100.00 | 100.00 | 100.00 |
| 3 | Brick | 6 | 4 | 0.30 | 332 | 84.44 | 250 | 332 | 364.58 | 460 | 77.78 | 82.65 | 84.44 |
| 4 | Mini Triangle | 6 | 4 | 0.30 | 30 | 100.00 | 250 | 30 | 30.00 | 30 | 100.00 | 100.00 | 100.00 |
| 5 | DeBruijn Sequence Square | 6 | 4 | 0.30 | 44 | 100.00 | 250 | 44 | 44.00 | 44 | 100.00 | 100.00 | 100.00 |
| 6 | puc19 Rectangle | 6 | 4 | 0.30 | 2 | 100.00 | 250 | 2 | 2.00 | 2 | 100.00 | 100.00 | 100.00 |
| 7 | 6 Helix Bundle | 6 | 4 | 0.30 | 238 | 93.75 | 250 | 201 | 260.76 | 396 | 89.58 | 93.39 | 94.79 |
| 8 | Fivewell Plate | 6 | 4 | 0.30 | 155 | 99.37 | 250 | 155 | 169.60 | 205 | 98.73 | 99.18 | 99.37 |
| 9 | Membrane Nanopore | 6 | 4 | 0.30 | – | – | – | – | – | – | – | – | – |
| 10 | Single Staple Loop | 6 | 4 | 0.30 | 2 | 100.00 | 250 | 2 | 2.00 | 2 | 100.00 | 100.00 | 100.00 |
| 11 | DNA Frame | 6 | 4 | 0.30 | 205 | 99.10 | 250 | 197 | 214.60 | 340 | 97.30 | 98.97 | 99.10 |
| 12 | Rothemund Rectangle | 6 | 4 | 0.30 | 126 | 100.00 | 250 | 126 | 126.00 | 126 | 100.00 | 100.00 | 100.00 |
| 13 | Rectangle Variant | 6 | 4 | 0.30 | 114 | 100.00 | 250 | 114 | 114.00 | 114 | 100.00 | 100.00 | 100.00 |
| 14 | Small Moon | 6 | 4 | 0.30 | 460 | 96.15 | 163 | 407 | 435.52 | 460 | 96.15 | 96.22 | 96.58 |
| 15 | Rothemund Smiley | 6 | 4 | 0.30 | 185 | 99.18 | 250 | 185 | 214.34 | 394 | 97.12 | 98.81 | 99.18 |
| 16 | Rothemund Star | 6 | 4 | 0.30 | 178 | 98.77 | 250 | 116 | 139.93 | 424 | 95.90 | 99.29 | 99.59 |
| 17 | Capsule | 6 | 4 | 0.30 | – | – | – | – | – | – | – | – | – |
| 18 | Triangle | 5 | 2 | 0.20 | 8 | 100.00 | 250 | 8 | 8.00 | 8 | 100.00 | 100.00 | 100.00 |
| 19 | Square | 5 | 2 | 0.20 | 6 | 100.00 | 250 | 6 | 6.00 | 6 | 100.00 | 100.00 | 100.00 |
| 20 | Pentagon | 5 | 2 | 0.20 | 6 | 100.00 | 250 | 6 | 6.00 | 6 | 100.00 | 100.00 | 100.00 |
| 21 | Tetrahedron | 5 | 2 | 0.20 | 4 | 100.00 | 250 | 4 | 4.00 | 4 | 100.00 | 100.00 | 100.00 |
| 22 | Triangle Mesh | 5 | 2 | 0.20 | 35 | 95.45 | 250 | 35 | 35.00 | 35 | 95.45 | 95.45 | 95.45 |
| 23 | Cube | 5 | 2 | 0.20 | 6 | 100.00 | 250 | 6 | 49.09 | 87 | 92.00 | 95.74 | 100.00 |
| 24 | Star Mesh | 5 | 2 | 0.20 | 66 | 97.37 | 250 | 66 | 66.00 | 66 | 97.37 | 97.37 | 97.37 |
| 25 | Dodecahedron | 5 | 2 | 0.20 | 188 | 96.61 | 250 | 188 | 226.30 | 272 | 93.22 | 95.06 | 96.61 |
| 26 | Icosahedron | 5 | 2 | 0.20 | 397 | 87.50 | 250 | 396 | 402.88 | 470 | 84.38 | 87.24 | 87.50 |
| 27 | Square Mesh 1 | 5 | 2 | 0.20 | 336 | 91.46 | 250 | 309 | 322.61 | 336 | 91.46 | 91.46 | 91.46 |
| 28 | Hexagon Mesh 1 | 5 | 2 | 0.20 | 358 | 91.67 | 250 | 333 | 369.62 | 528 | 85.71 | 90.81 | 92.86 |
| 29 | Annulus Mesh 1 | 5 | 2 | 0.20 | 91 | 98.88 | 250 | 91 | 91.00 | 91 | 98.88 | 98.88 | 98.88 |
| 30 | Hexagonal Tile | 5 | 2 | 0.20 | 169 | 97.96 | 250 | 169 | 169.00 | 169 | 97.96 | 97.96 | 97.96 |
| 31 | Truncated Cube | 5 | 2 | 0.20 | 72 | 100.00 | 250 | 72 | 72.00 | 72 | 100.00 | 100.00 | 100.00 |
| 32 | Cross Mesh | 5 | 2 | 0.20 | 226 | 96.36 | 250 | 226 | 256.15 | 340 | 94.55 | 96.10 | 96.36 |
| 33 | vHelix Ball | 5 | 2 | 0.20 | 18 | 100.00 | 250 | 18 | 18.00 | 18 | 100.00 | 100.00 | 100.00 |
| 34 | Annulus Mesh 2 | 5 | 2 | 0.20 | 154 | 99.38 | 250 | 142 | 173.15 | 202 | 98.76 | 99.06 | 99.38 |
| 35 | Lotus Mesh | 5 | 2 | 0.20 | 302 | 97.78 | 250 | 273 | 287.50 | 302 | 97.78 | 97.78 | 97.78 |
| 36 | Enneagonal Trapezohedron | 5 | 2 | 0.20 | – | – | – | – | – | – | – | – | – |

Supplementary Table 2: REVNANO run with consensus parameters in deterministic and non-deterministic mode. Column $n$ is the number of non-deterministic repeats over which statistics are calculated (i.e. that did not end with a REVNANO error). Values highlighted in blue are cases when non-deterministic mode yields better results (over $n$ repeats) than deterministic mode.

| | | Deterministic | Best non-deterministic | | Deterministic | Best non-deterministic | |
|---|---|---|---|---|---|---|---|
| | | | over multiple samples | | | over multiple samples | |
| ID | Origami | $d(\mathbf{g}, \mathbf{r})$ | min $d(\mathbf{g}, \mathbf{r})$ | Diff | SP% | max SP% | Diff |
| 7 | 6 Helix Bundle | 238 | 201 | -37 | 93.75 | 94.79 | +1.04 |
| 11 | DNA Frame | 205 | 197 | -8 | 99.10 | 99.10 | 0 |
| 14 | Small Moon | 460 | 407 | -53 | 96.15 | 96.58 | +0.43 |
| 16 | Rothemund Star | 178 | 116 | -62 | 98.77 | 99.59 | +0.82 |
| 26 | Icosahedron | 397 | 396 | -1 | 87.50 | 87.50 | 0 |
| 27 | Square Mesh 1 | 336 | 309 | -27 | 91.46 | 91.46 | 0 |
| 28 | Hexagon Mesh 1 | 358 | 333 | -25 | 91.67 | 92.86 | +1.19 |
| 34 | Annulus Mesh 2 | 154 | 142 | -12 | 99.38 | 99.38 | 0 |
| 35 | Lotus Mesh | 302 | 273 | -29 | 97.78 | 97.78 | 0 |

Supplementary Table 3: Analysis of origamis from Supplementary Table 2 where non-deterministic staple placement (over multiple repeats) can cause better REVNANO performance than deterministic mode. Improvements are marginal.

### Supplementary Note 5 REVNANO with Noisy Staple Sequences

The REVNANO solver critically depends on finding contiguous complementary Watson-Crick regions between staples and the scaffold strand. It assumes that all staples are present, and that no staple sequence errors exist. The following numerical analysis investigates how quickly REVNANO performance degrades when various types of 'noise' are added to the staple sequences input into the solver.

Noise may result when staple sequences are not copied from a publication correctly, when unrelated staple sequences are included by accident, or when published sequences themselves are incomplete and/or contain base errors.

A 2D raster- and a 3D wireframe shape were tested with three different types of staple noise detailed in Supplementary Figure 4. For both origamis, REVNANO was run with consensus parameters and in deterministic mode. Noise was not added to the scaffold sequence: it is assumed that the scaffold sequence is copied from a standard sequence repository and is correct.

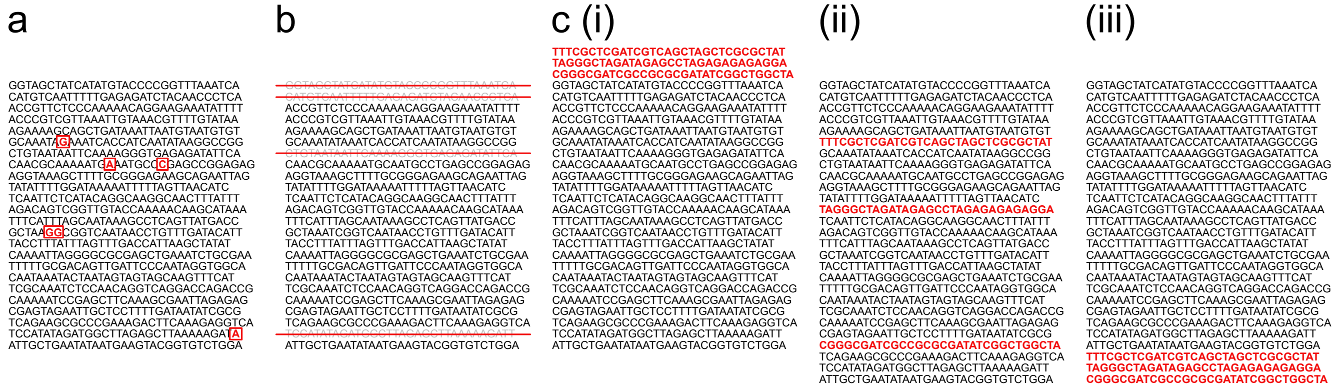


Supplementary Figure 4: Three types of staple noise investigated. The correct full staple list of the origami has: (a) single staple bases randomly perturbed such that they become non-complementary to the scaffold strand; (b) randomly selected staples omitted; (c) extra unrelated random sequence DNA staples (not part of the origami) inserted at the beginning (i), randomly (ii) or at the end (iii). In case (b), omitted staples are randomly selected without replacement. In case (c), extra staples have length assigned randomly between the shortest and longest staples in the correct full staple list.

Here we report on how two metrics change as staple noise increases. The first metric is the hamming distance $d(\mathbf{g}, \mathbf{r})$ from the ground truth contact map $\mathbf{r}$ to the REVNANO reconstructed contact map $\mathbf{g}$ for the origami shape, in the presence of noise. Hamming distance error generally increases as noise increases. We note that REVNANO contact maps reconstructed in noise conditions (b), c(i) and c(ii) of Supplementary Figure 4 required staples to be re-numbered to have the same ID numbers as they did in the original ground truth contact map, in order for hamming distance to be calculated meaningfully.

The second metric is surviving staples fraction:

$$\text{SS\%} = \frac{\text{number of staples in ground truth contact map placed in reconstructed REVNANO contact map}}{\text{total number of staples in ground truth contact map}} \tag{1}$$

which reports what fraction of staples in the original correct staple set are placed in the reconstructed REVNANO contact map, when noise is present. Importantly, this metric definition also allows to deal with cases where noise is manifest as extra unrelated staples (Supplementary Figure 4c). Surviving staples fraction generally decreases as noise increases.

Supplementary Figures 5, 6, 7 and 8 below reveal two main results:

1. The REVNANO solver is most sensitive to staple noise in the form of perturbed staple bases, or staples which are omitted (Supplementary Figures 5 and 7). These two types of noise have approximately equal detrimental effects on REVNANO performance. As more staple bases are perturbed (or more staples are omitted) the hamming distance error monotonically increases (with some variance). When noise comes in the form of perturbed staple bases, hamming distance error starts to level off at high noise levels, and a point comes when REVNANO suddenly cannot produce a solution (red arrows on figures). On the other hand, when noise is in the form of omitted staples, hamming distance error is linear in number of staples omitted.

2. When extra random DNA staples are inserted at the end of the correct staple list (part c(iii) of Supplementary Figures 6 and 8), in many cases REVNANO can still reconstruct an accurate contact map, effectively 'fishing' the correct staple sequences out of a contaminated staple sequence pool (in a minority of cases, however, the extra staples have chance DNA sequences which significantly interfere with contact map reconstruction). On the other hand, inserting extra random DNA staples at the start of the correct staple list is more detrimental to REVNANO performance, and the correct nanostructure can be 'fished out' more infrequently. Inserting extra staples at random positions in the staple list has a performance hit between the above two extremes. REVNANO robustness to extra DNA staples added at the end of the correct staple can be expected, considering that in deterministic mode these staples are polled last (after the preceding 'correct' staples have already established their constraints).

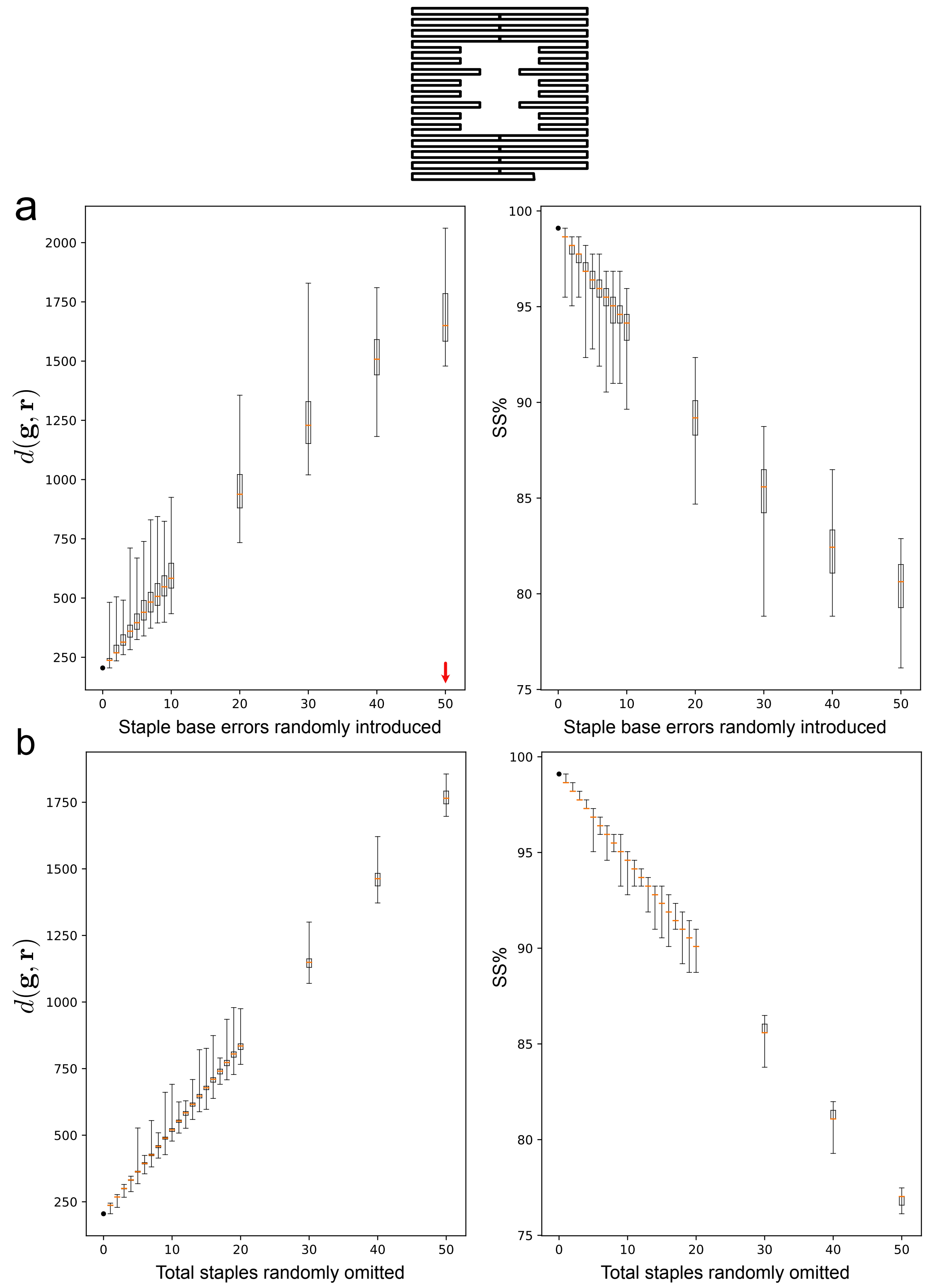


Supplementary Figure 5: Origami 11 (DNA Frame): 7249nt scaffold, 222 staples, 7112 staple bases hybridised to scaffold. Box plots showing REVNANO reverse engineering performance using consensus parameters ($\mu_{\min} = 6\text{bp}, \sigma = 4\text{bp}, \beta = 0.3$) in presence of (a) increasing single base errors on staples, (b) increasing number of staples omitted. Orange lines indicate median values. Red arrows denote no further solutions beyond this parameter value. See Supplementary Table 4 for REVNANO repeats performed at each parameter point.

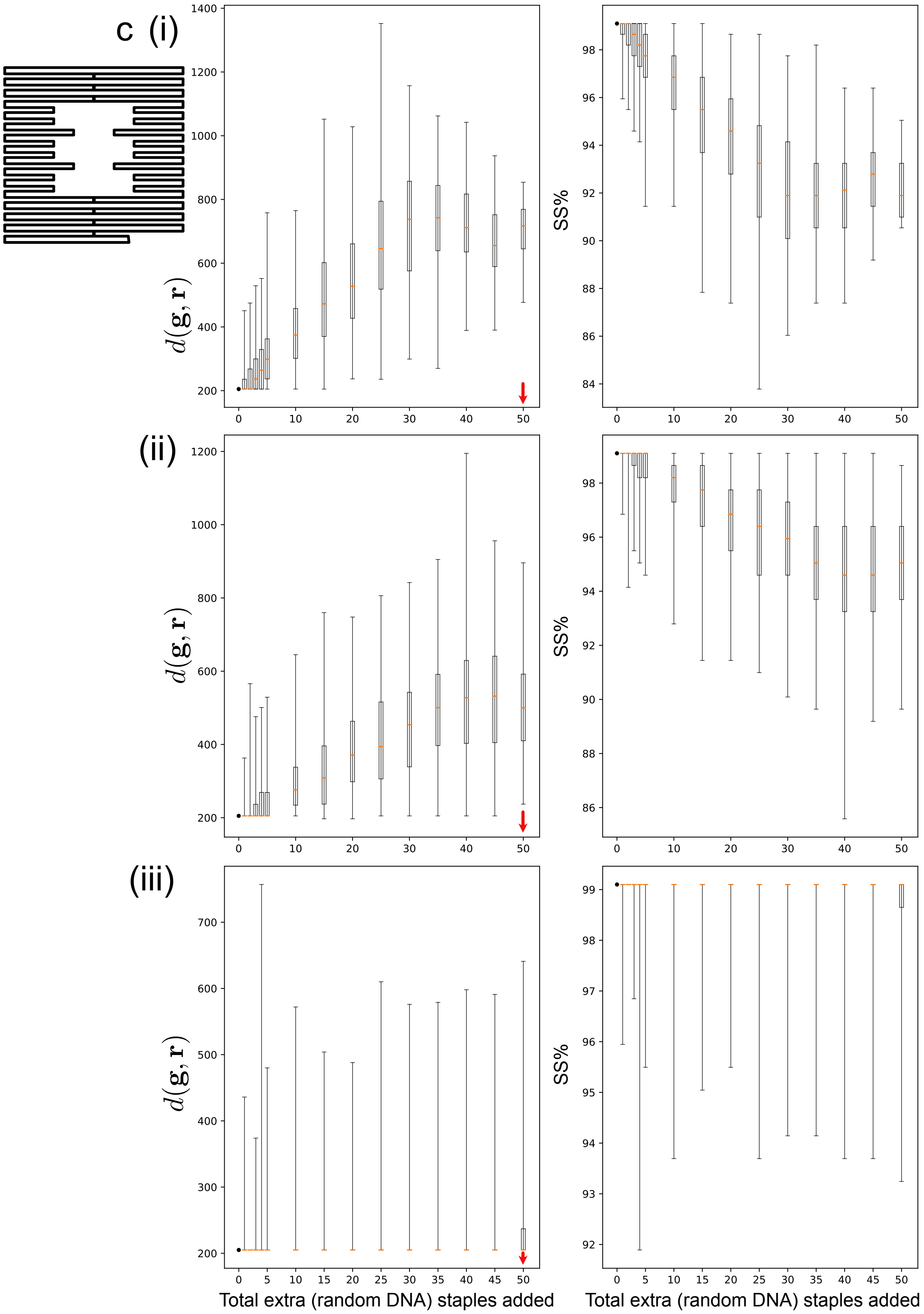


Supplementary Figure 6: Origami 11 Continued from Supplementary Figure 5. Box plots showing REVNANO reverse engineering performance in presence of (c) increasing numbers of unrelated random DNA staples not in the origami design, inserted at (i) beginning, (ii) random locations, (iii) end of staple list. Orange lines indicate median values. Red arrows denote no further solutions beyond this parameter value. See Supplementary Table 4 for REVNANO repeats performed at each parameter point.

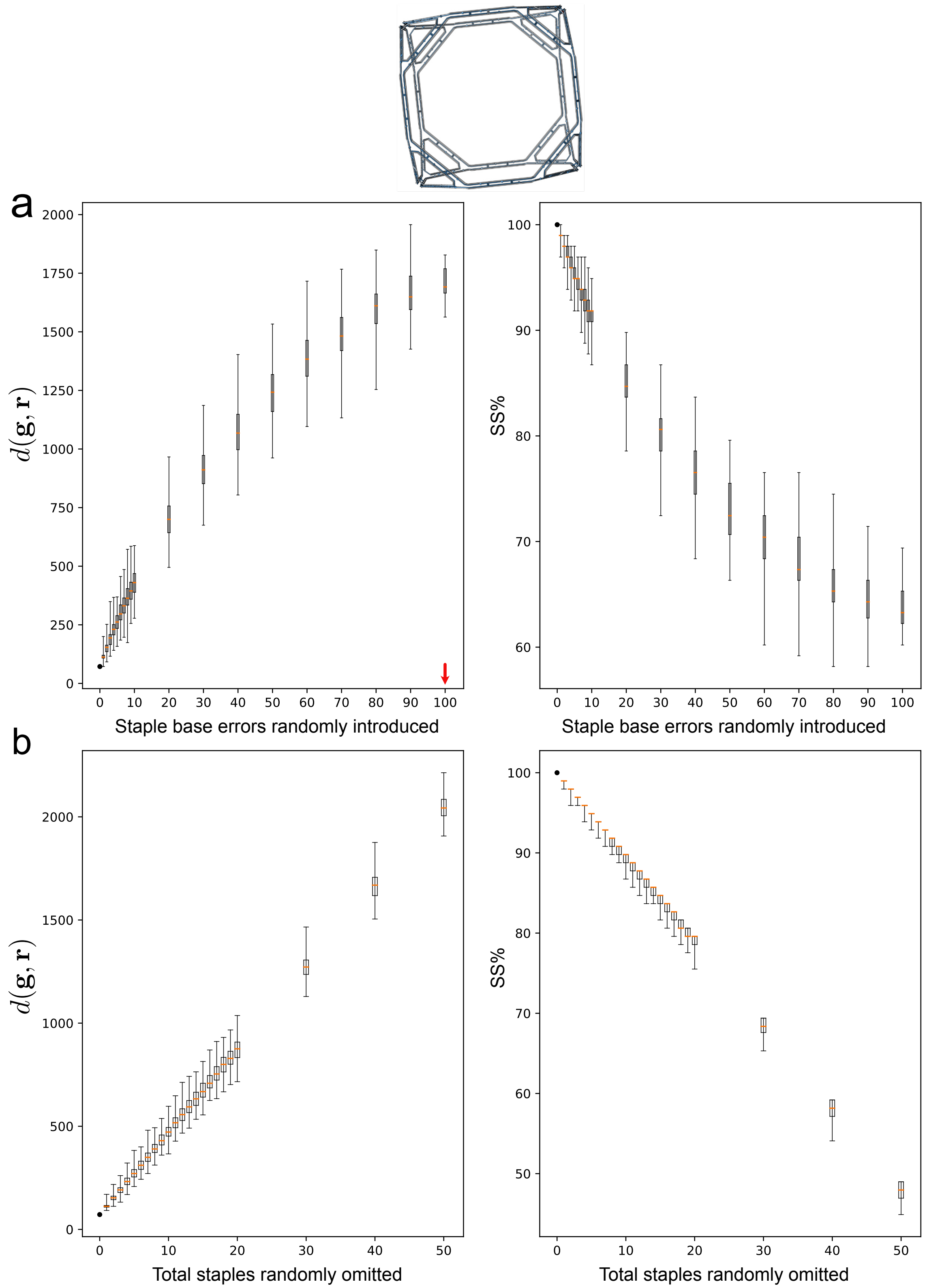


Supplementary Figure 7: Origami 31 (Truncated Cube): 3960nt scaffold, 98 staples, 3840 staple bases hybridised to scaffold. Box plots showing REVNANO reverse engineering performance using consensus parameters $(\mu_{\min} = 5\mathrm{bp}, \sigma = 2\mathrm{bp}, \beta = 0.2)$ in presence of (a) increasing single base errors on staples, (b) increasing number of staples omitted. Orange lines indicate median values. Red arrows denote no further solutions beyond this parameter value. See Supplementary Table 4 for REVNANO repeats performed at each parameter point.

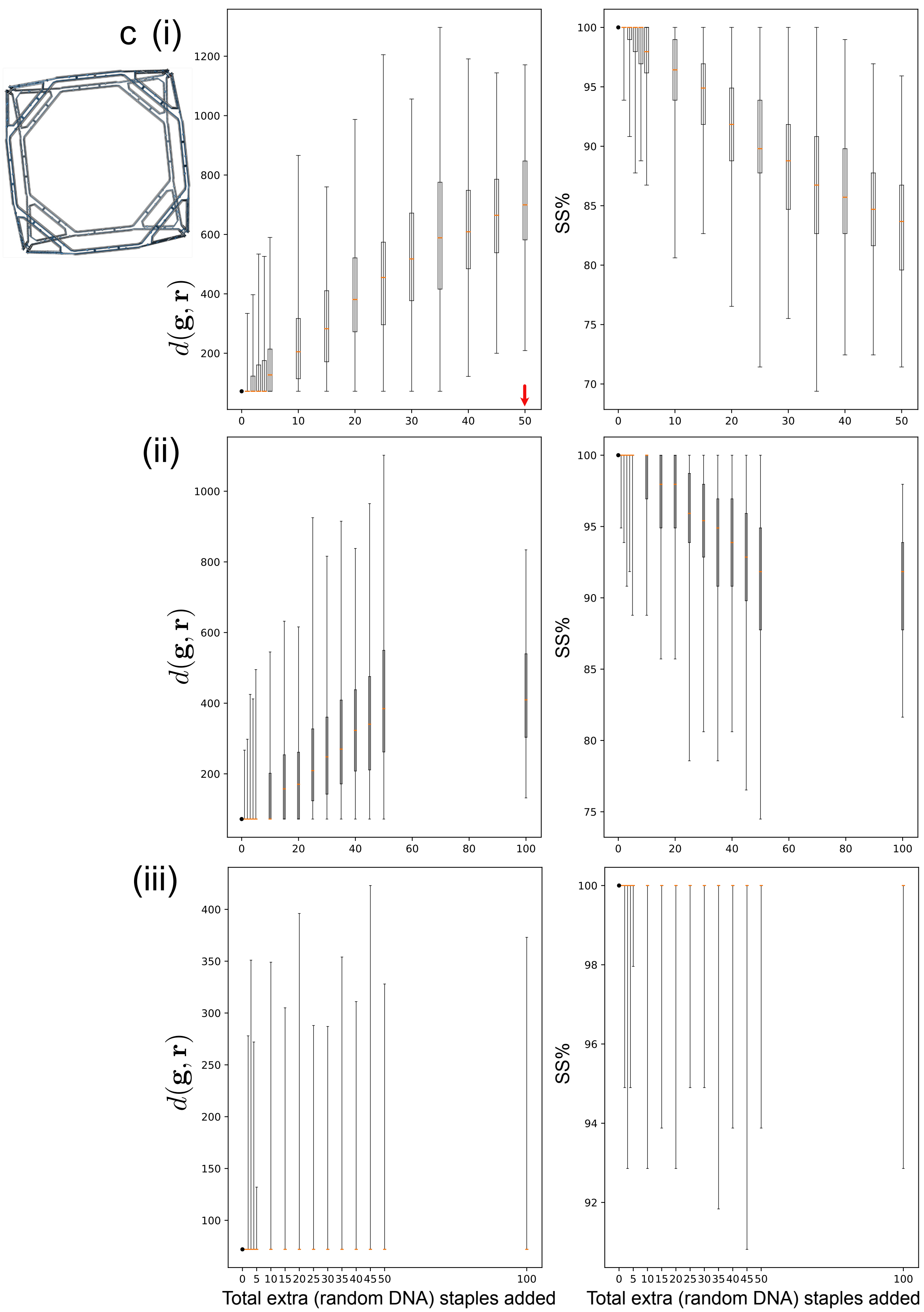


Supplementary Figure 8: Origami 31 Continued from Supplementary Figure 7. Box plots showing REVNANO reverse engineering performance in presence of (c) increasing numbers of unrelated random DNA staples not in the origami design, inserted at (i) beginning, (ii) random locations, (iii) end of staple list. Orange lines indicate median values. Red arrows denote no further solutions beyond this parameter value. See Supplementary Table 4 for REVNANO repeats performed at each parameter point.

| | 5a | 7a | | 5b | 7b | | 6c(i) | 6c(ii) | 6c(iii) | 8c(i) | 8c(ii) | 8c(iii) |
|---|---|---|---|---|---|---|---|---|---|---|---|---|
| Parameter | $n$ | $n$ | Parameter | $n$ | $n$ | Parameter | $n$ | $n$ | $n$ | $n$ | $n$ | $n$ |
| 1 | 250 | 250 | 1 | 250 | 250 | 1 | 250 | 250 | 250 | 250 | 250 | 250 |
| 2 | 250 | 250 | 2 | 250 | 250 | 2 | 250 | 250 | 250 | 250 | 250 | 250 |
| 3 | 250 | 250 | 3 | 250 | 250 | 3 | 250 | 250 | 250 | 250 | 250 | 250 |
| 4 | 250 | 250 | 4 | 250 | 250 | 4 | 250 | 250 | 250 | 250 | 250 | 250 |
| 5 | 250 | 250 | 5 | 250 | 250 | 5 | 250 | 250 | 250 | 250 | 250 | 250 |
| 6 | 250 | 250 | 6 | 250 | 250 | 10 | 250 | 250 | 250 | 250 | 250 | 250 |
| 7 | 250 | 250 | 7 | 250 | 250 | 15 | 250 | 250 | 250 | 250 | 250 | 250 |
| 8 | 250 | 250 | 8 | 250 | 250 | 20 | 250 | 250 | 250 | 250 | 250 | 250 |
| 9 | 250 | 250 | 9 | 250 | 250 | 25 | 247 | 250 | 250 | 249 | 250 | 250 |
| 10 | 250 | 250 | 10 | 250 | 250 | 30 | 213 | 247 | 250 | 250 | 250 | 250 |
| 20 | 250 | 250 | 11 | 250 | 250 | 35 | 166 | 246 | 250 | 250 | 250 | 250 |
| 30 | 225 | 250 | 12 | 250 | 250 | 40 | 98 | 234 | 250 | 247 | 250 | 250 |
| 40 | 85 | 249 | 13 | 250 | 250 | 45 | 46 | 197 | 250 | 242 | 250 | 250 |
| 50 | 13 | 250 | 14 | 250 | 250 | 50 | 13 | 141 | 250 | 228 | 250 | 250 |
| 60 | 1 | 250 | 15 | 250 | 250 | 100 | 0 | 0 | 0 | 3 | 34 | 246 |
| 70 | 0 | 237 | 16 | 250 | 250 | | | | | | | |
| 80 | 0 | 190 | 17 | 250 | 250 | | | | | | | |
| 90 | 0 | 99 | 18 | 250 | 250 | | | | | | | |
| 100 | 0 | 28 | 19 | 250 | 250 | | | | | | | |
| 200 | 0 | 0 | 20 | 250 | 250 | | | | | | | |
| | | | 30 | 249 | 250 | | | | | | | |
| | | | 40 | 211 | 250 | | | | | | | |
| | | | 50 | 110 | 250 | | | | | | | |

Supplementary Table 4: Sample sizes for box plots in Supplementary Figures 5, 6, 7 and 8. Figure numbers indicated in top row. Every x-axis parameter point (x>0) in the latter figures ideally executes REVNANO $n = 250$ times with a different random seed. However, 250 samples are not always possible to perform as REVNANO can terminate with an error or sometimes does not terminate at all in reasonable time. Thus, actual sample sizes for all box plots are listed above. A box plot is only drawn when 10 samples or more exist.

### Supplementary Note 6 REVNANO with Staple Dangles and Staple Loopouts

Staple dangles and loopouts are quite common features in practical origamis:

- Staple dangles are used at origami edges to reduce or enhance origami-origami stacking, or used as functional protruding sequences to e.g. secure other DNA or RNA strands to an origami.
- Staple loopouts are used extensively in 2D and 3D wireframe origamis to make various types of junction.
- 24 of 36 origamis reported in the paper have a percentage of staples with either end dangles and/or interior loopouts.

REVNANO relies on the fact that all bases of a staple hybridise with the origami scaffold. The algorithm attempts to find number and positions of all crossovers on each staple. Regions of staple sequences which are (i) single-stranded end dangles or (ii) single-stranded interior loopouts (see Supplementary Figure 1a) must be manually enclosed with * pairs in the REVNANO input file, such that the algorithm knows to ignore them.

The REVNANO algorithm cannot automatically decide where non-hybridising regions are located on a staple (if any exist at all) because this feature expands the search space of the core constraint satisfaction problem significantly: there exists a combinatorially large number of ways such non-hybridising regions could appear on each staple (in terms of locations and lengths). Also, rather than one unique solution, multiple optimal solutions of the CSP would likely exist.

The best option for using the REVNANO solver with origamis containing staples with dangles and/or interior loopouts ("DL" staples) is to use information in the original paper and supplementary information to manually mark these sequence regions in the REVNANO input file. If no information at all exists about the number or locations of dangles and/or interior loopout regions on DL staples, then two options remain:

1. The REVNANO solver can be tried as a *detector* of DL staples. Ideally, the staples that REVNANO is unable to fully route are precisely the DL staples. The user can review these staples in conjunction with the guide schematic, in order to infer the non-hybridising regions and manually mark them.
2. Non-hybridising regions are usually polyT: all polyT regions (i.e. TTTT or longer) of all staple sequences can be enclosed with * pairs, even though some of these marked polyT regions will actually hybridise with the scaffold in the origami design. Ideally, this will protect the bona fide dangle and/or loopout regions in the staples from REVNANO, but it will also protect many other staples regions which do in fact hybridise with the scaffold. Ideally, however, the user can make use of the guide schematic to rectify the dangle/loopout regions which are incorrectly identified.

Supplementary Table 5 shows Option 1 results. All origamis with DL staples have a REVNANO input file constructed where dangle and/or interior loopout regions are not marked. The results show that REVNANO can indeed be used as a detector for DL staples, but this only works in limited cases, i.e. for Origami 7 and 19 where the staples omitted are precisely the set of DL staples. In most cases however, for the parameters tried, the solver will (incorrectly) incorporate some DL staples into the origami. This leads to disruption in the proper staple routing and in many cases causes the solver to terminate in an error.

| | | | | | | DL Marked | | DL Not Marked | | | |
|---|---|---|---|---|---|---|---|---|---|---|---|
| ID | Origami | Staples | DL Staples | Total Dangles | Total Loopouts | $d(\mathbf{g},\mathbf{r})$ | SP% | $d(\mathbf{g},\mathbf{r})$ | SP% | DL Incorporated | DL Omitted |
| 3 | Brick | 45 | 24 | 36 | 0 | 332 | 84.44 | REVNANO Error | | | |
| 4 | Mini Triangle | 66 | 9 | 0 | 9 | 30 | 100.00 | 416 | 84.85 | 5 | 4 |
| 6 | puc19 Rectangle | 90 | 12 | 24 | 0 | 2 | 100.00 | 380 | 86.67 | 0 | 12 |
| 7 | 6 Helix Bundle | 96 | 8 | 12 | 0 | 136 | 96.88 | 443 | 86.46 | 1 | 7 |
| 13 | Rectangle Variant | 226 | 24 | 4 | 22 | 114 | 100.00 | 1006 | 88.50 | 8 | 16 |
| 18 | Triangle | 7 | 4 | 0 | 6 | 8 | 100.00 | REVNANO Error | | | |
| 19 | Square | 8 | 4 | 0 | 8 | 6 | 100.00 | 215 | 50.00 | 0 | 4 |
| 20 | Pentagon | 10 | 5 | 0 | 10 | 6 | 100.00 | REVNANO Error | | | |
| 21 | Tetrahedron | 13 | 7 | 0 | 12 | 4 | 100.00 | 287 | 46.15 | 1 | 6 |
| 22 | Triangle Mesh | 22 | 13 | 0 | 18 | 35 | 95.45 | REVNANO Error | | | |
| 23 | Cube | 25 | 13 | 0 | 24 | 6 | 100.00 | REVNANO Error | | | |
| 24 | Star Mesh | 38 | 20 | 0 | 36 | 66 | 97.37 | REVNANO Error | | | |
| 25 | Dodecahedron | 59 | 30 | 0 | 60 | 188 | 96.61 | REVNANO Error | | | |
| 26 | Icosahedron | 64 | 41 | 0 | 60 | 355 | 89.06 | REVNANO Error | | | |
| 27 | Square Mesh 1 | 82 | 42 | 0 | 62 | 284 | 92.68 | REVNANO Error | | | |
| 28 | Hexagon Mesh 1 | 84 | 49 | 0 | 72 | 335 | 92.86 | REVNANO Error | | | |
| 29 | Annulus Mesh 1 | 89 | 41 | 0 | 60 | 91 | 98.88 | REVNANO Error | | | |
| 30 | Hexagonal Tile | 98 | 49 | 0 | 98 | 169 | 97.96 | REVNANO Error | | | |
| 31 | Truncated Cube | 98 | 62 | 0 | 72 | 72 | 100.00 | REVNANO Error | | | |
| 32 | Cross Mesh | 110 | 62 | 0 | 100 | 176 | 97.27 | REVNANO Error | | | |
| 34 | Annulus Mesh 2 | 161 | 76 | 0 | 100 | 106 | 100.00 | REVNANO Error | | | |
| 35 | Lotus Mesh | 180 | 66 | 0 | 93 | 271 | 97.78 | REVNANO Error | | | |

Supplementary Table 5: REVNANO used as a detector of DL staples. Only origamis with staple dangles and loopouts are listed. Origamis 17 and 36 are omitted. REVNANO is run with optimal parameters for each shape, listed in Table 1 of the paper. 'Total Dangles' are the total number of dangling ends present across all staples (similarly for 'Total Loopouts'). 'DL Incorporated' is the number of DL staples that REVNANO incorrectly routes in the origami, hybridising all bases. 'DL Omitted' is the number of DL staples that REVNANO was unable to route. Finally, REVNANO only serves as a DL staple detector for Origamis 7 and 19 (blue).

The alternative Option 2 of marking all polyT regions in all staples has the advantage that REVNANO can often recover some form of contact map for an origami (data not shown). However, often the "noise" of the incorrectly identified single-stranded staple regions outweighs the "signal" of the correctly identified single-stranded staple regions: this results in a guide schematic which is a tangled mess of connections, not resembling an origami shape. Such an unstructured guide schematic cannot be used to decipher which single-stranded staple regions should be dropped from the input file.

Overall, staple dangle and interior loopout regions present a non-trivial challenge to the REVNANO solver and are best dealt with by manually marking them in the REVNANO input file.

### Supplementary Note 7 REVNANO Parameter Space Examples

Full REVNANO parameter spaces for the Smiley Face and Annulus Mesh 2 origamis are displayed below, calculated by High Performance Computing. REVNANO is run in deterministic mode.

The optimal parameter point is defined as the lowest value of $\sigma$ with the minimum base hamming distance, and the lowest value of $\beta$ for the latter value of $\sigma$. Lower $\sigma$ leads to faster REVNANO computation times as staple routing trees are smaller.

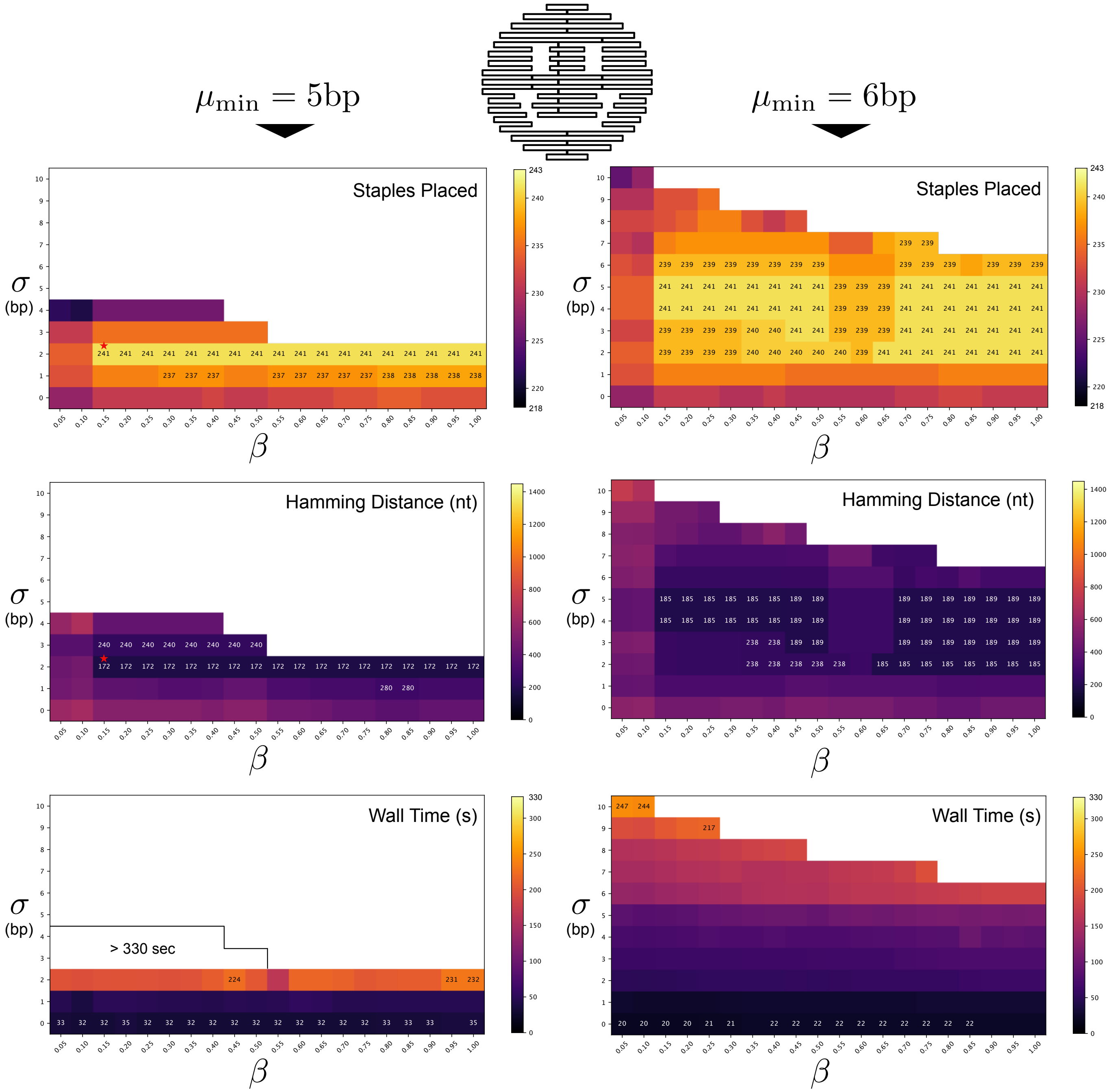


Supplementary Figure 9: Complete REVNANO parameter space for Rothemund Smiley (Origami 15). Left: $\mu_{\min} = 5$bp, Right: $\mu_{\min} = 6$bp. Top row heatmaps show total staples placed. Labelled squares show three highest values. Middle row heatmaps show base hamming distance to ground truth contact map $d(\mathbf{g}, \mathbf{r})$. Labelled squares show three lowest values. Bottom row heatmaps show execution time (in seconds) of each parameter point. Labelled squares show three highest and three lowest values. Squares appear white if data value is outside the colour bar range, or if REVNANO terminates with error. Red star marks optimal parameter point.

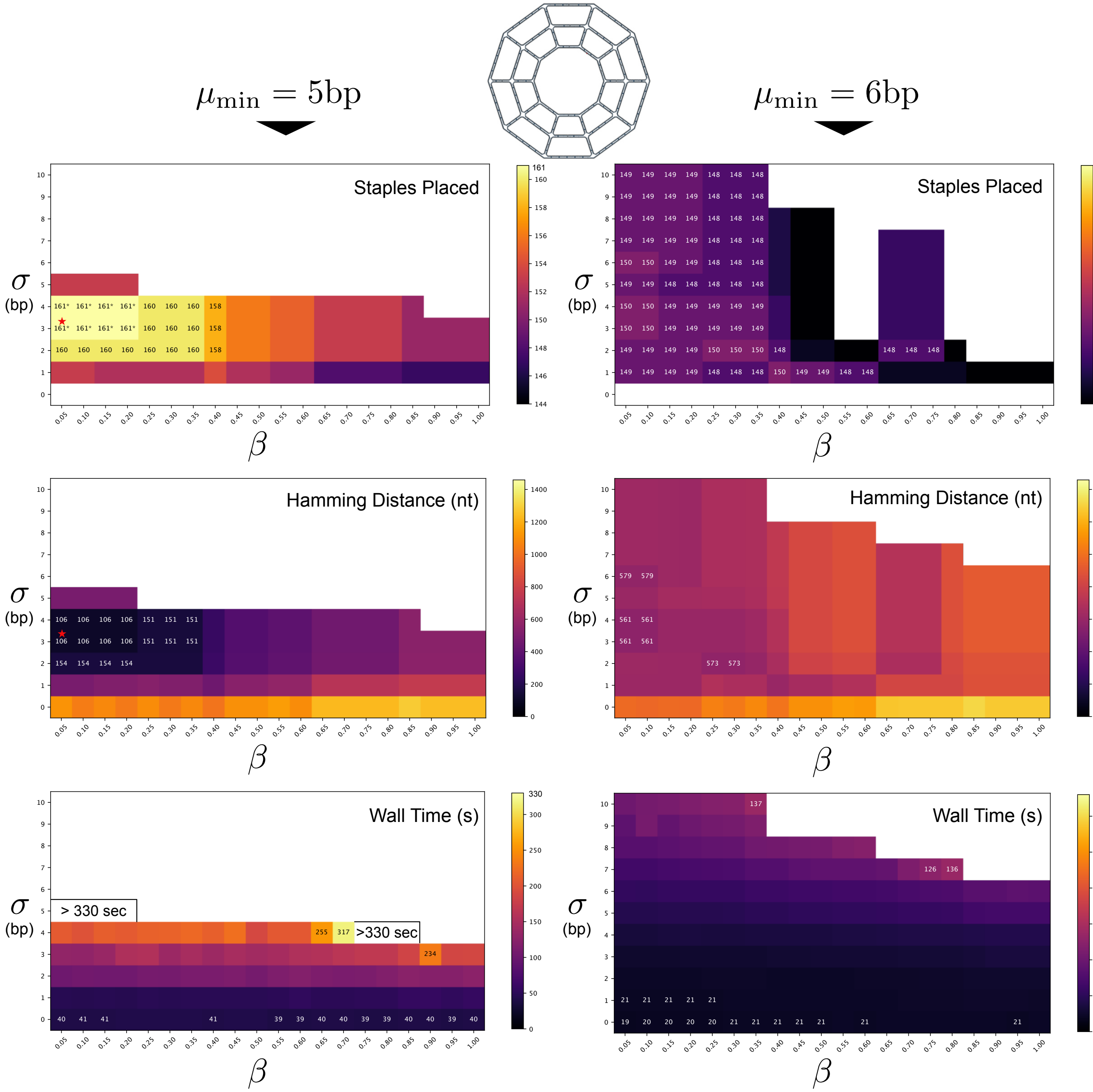


Supplementary Figure 10: Complete REVNANO parameter space for Annulus Mesh 2 (Origami 34). Left: $\mu_{\min} = 5$bp, Right: $\mu_{\min} = 6$bp. Top row heatmaps show total staples placed. Labelled squares show three highest values. Middle row heatmaps show base hamming distance to ground truth contact map $d(\mathbf{g}, \mathbf{r})$. Labelled squares show three lowest values. Bottom row heatmaps show execution time (in seconds) of each parameter point. Labelled squares show three highest and three lowest values. Squares appear white if data value is outside the colour bar range or if REVNANO terminates with error. Red star marks optimal parameter point.

#### Supplementary Note 8 Reverse Engineered Guide Schematics

The following pages display guide schematics reverse engineered from staple/scaffold sequences, for all origamis in Table 1 of the paper (omitting those already presented in Figure 4 of the paper). In each case, REVNANO was used with optimal parameters listed in Table 1 of the paper.

Origamis with red numbers indicate that the guide schematic is not interpretable (due to bad layout and/or dense internal connectivity). Five such schematics exist (Origamis 3, 7, 9, 27, 28). Origamis with starred numbers indicate that the guide schematic has unhybridised 1nt domains omitted from the scaffold strand (see Supplementary Note 1).

Thumbnail schematics for 2D raster origamis were exported using scadnano[1]. Thumbnails for 3D raster origamis were exported using CanDo[5, 6, 7] and UCSF Chimera[8]. Thumbnail schematics for wireframe origamis were produced by ATHENA[4] and exported by UCSF Chimera[8].

Supplementary Figure 11 highlights the interactive features of the guide schematic HTML page, using Origami 32.

Supplementary Figure 12 shows the five origamis with poor guide schematics re-rendered from perfect ground truth contact maps (to show the hypothetical case of REVNANO reaching 100% staples placed). The guide schematic for two of these origamis (Origamis 27 and 28) becomes useful/interpretable.

Supplementary Figure 13 displays Origamis 17 and 36 (whose contact maps could not be solved by REVNANO) rendered from their ground truth contact maps to show the hypothetical case of REVNANO reaching 100% staples placed.

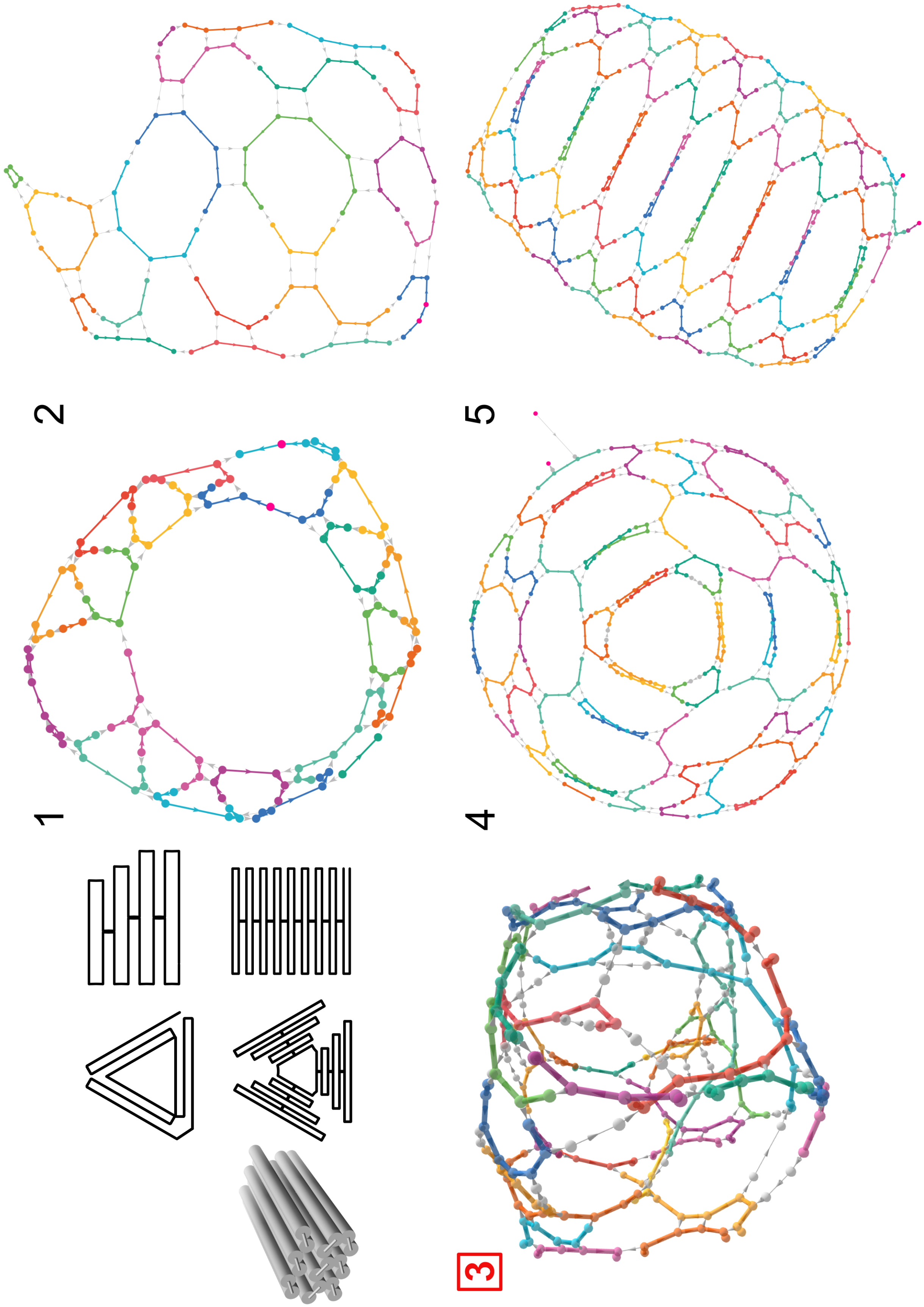
1
2
3
4
5

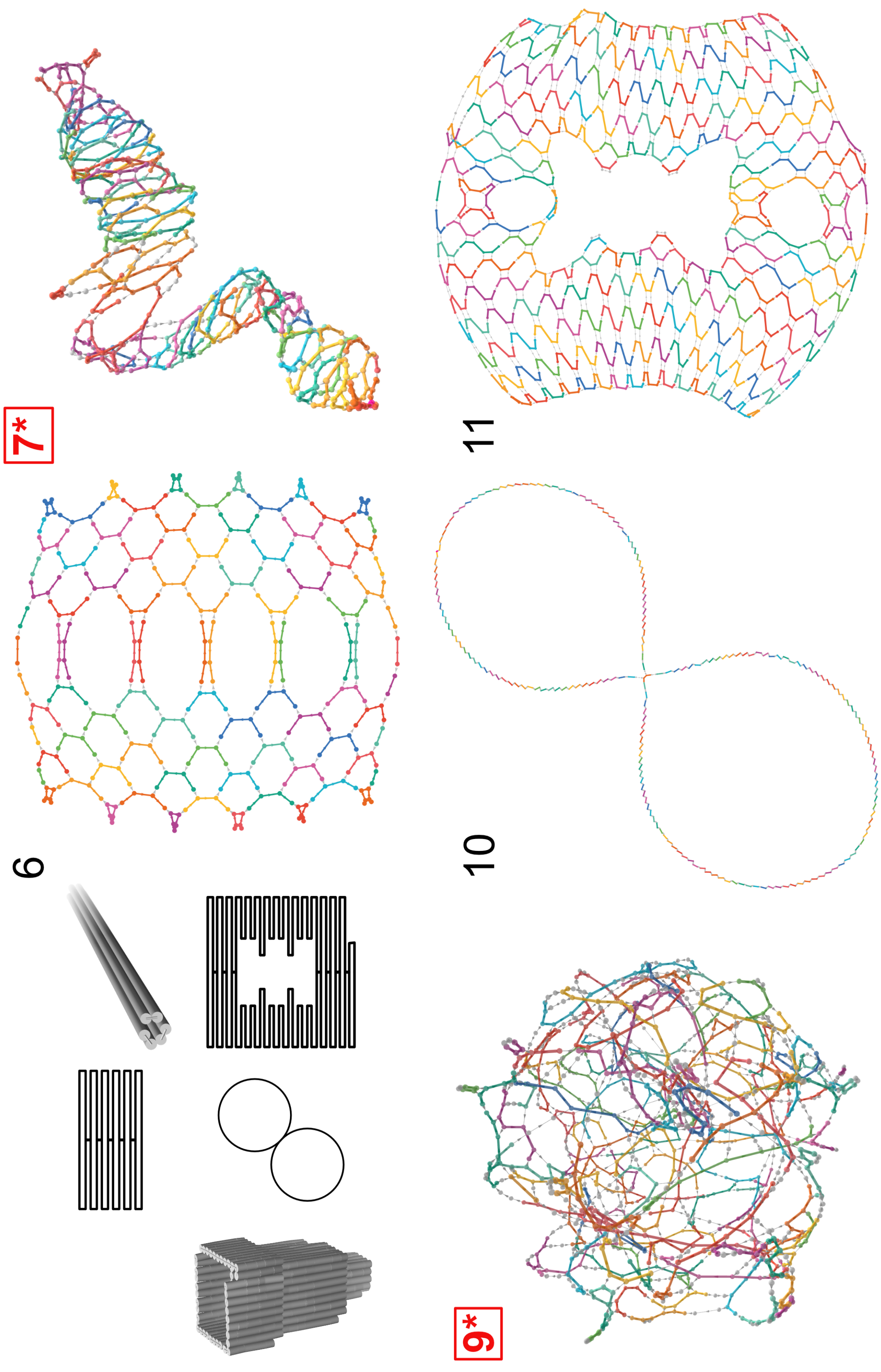
7*
6
11
10
9*

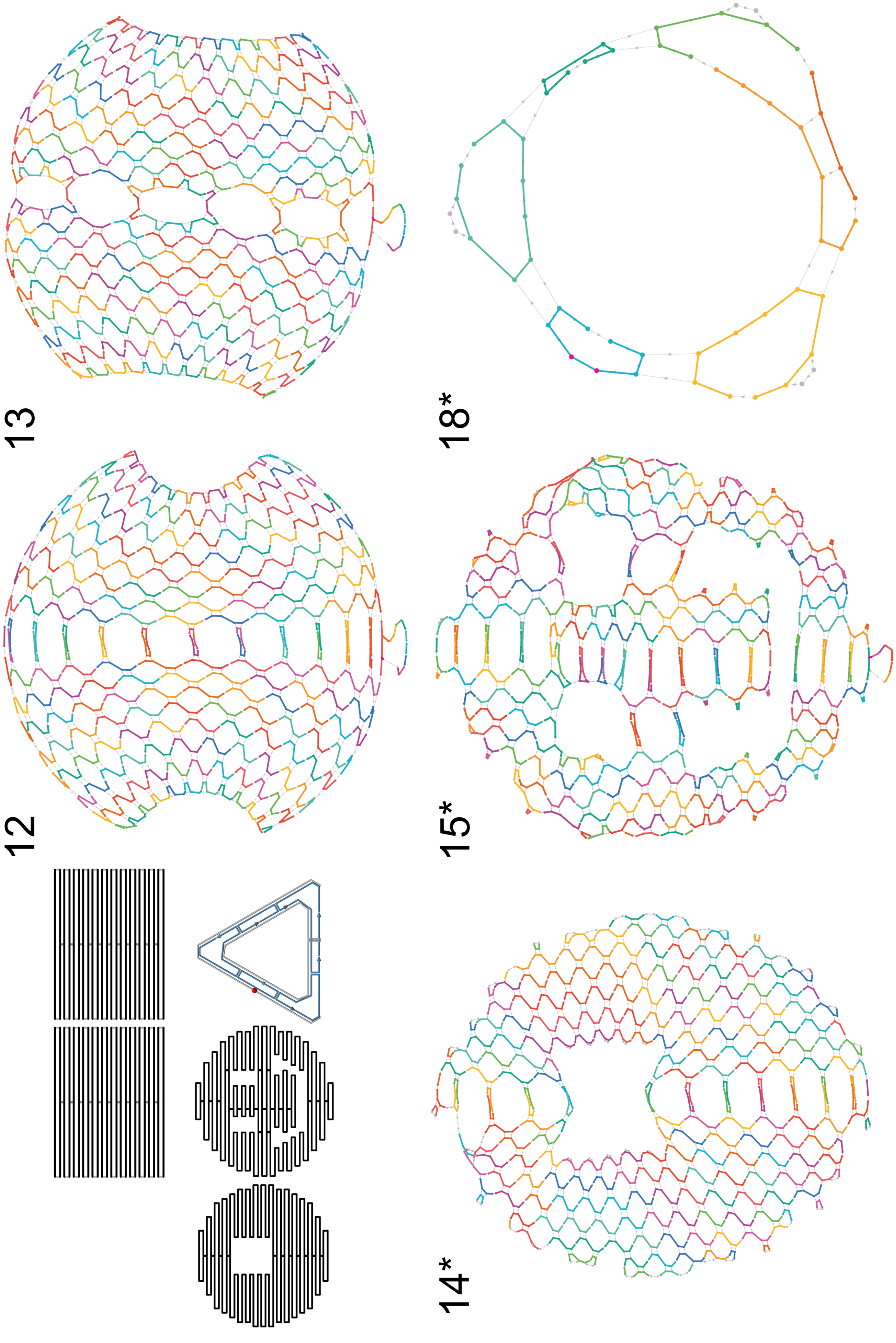
12
13
14*
15*
18*

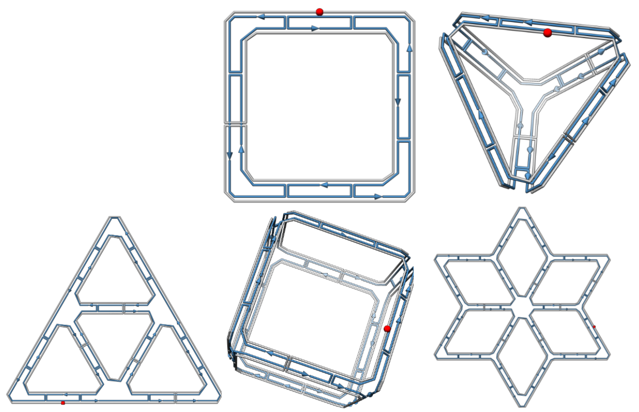

19*

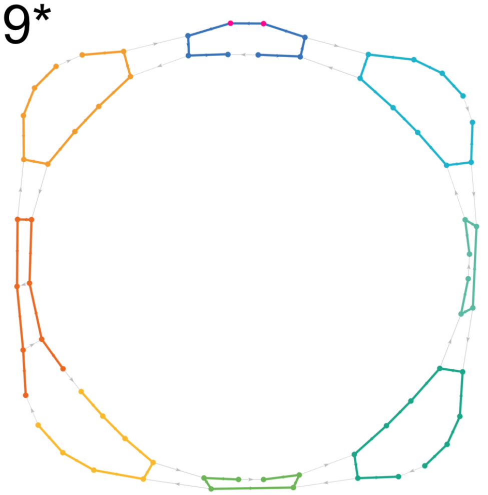

21

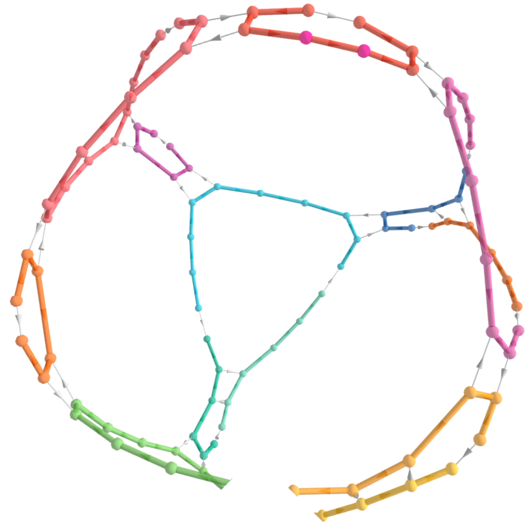

22*

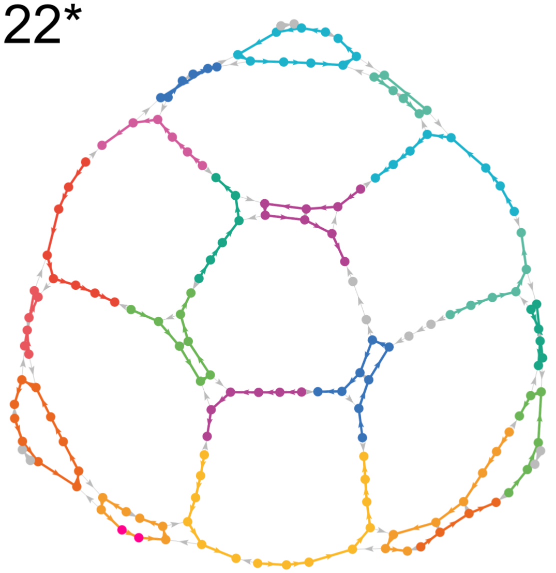

23

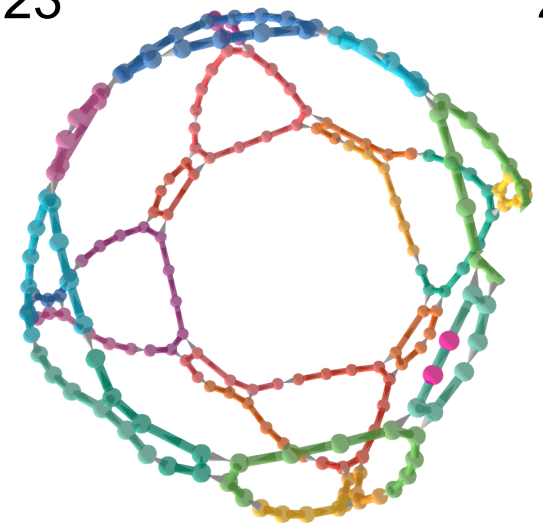

24*

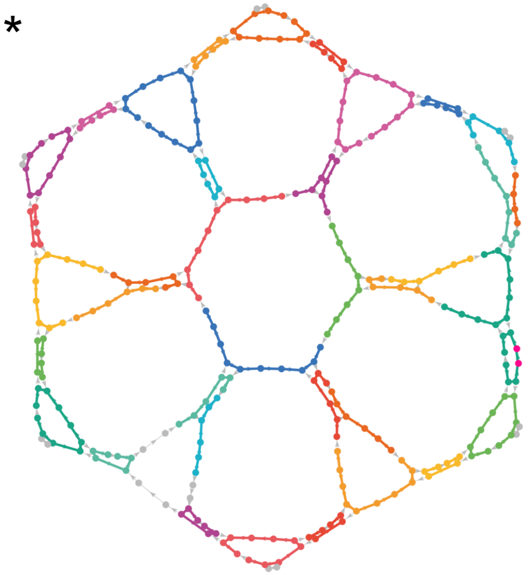

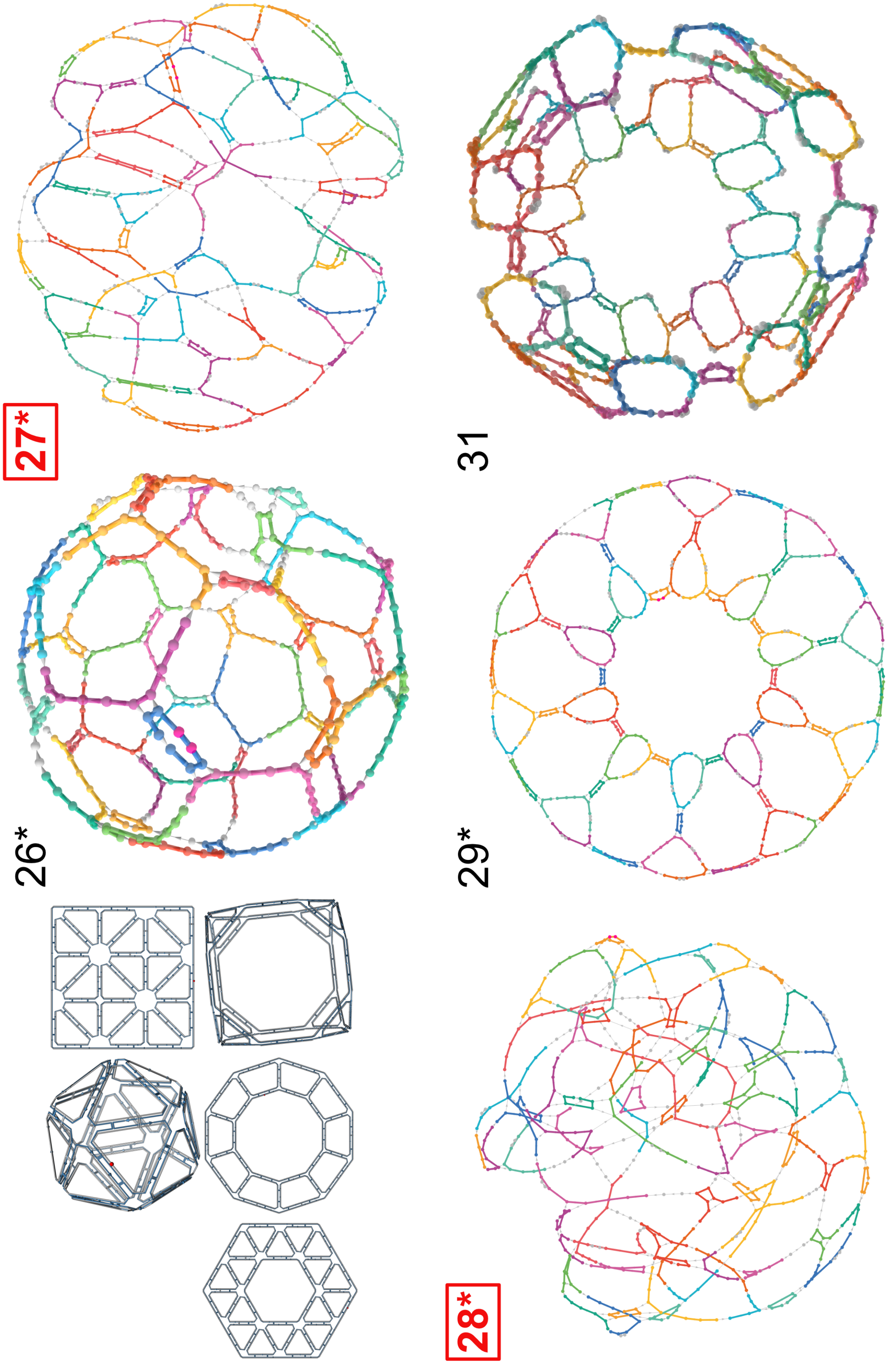
26*
27*
28*
29*
31

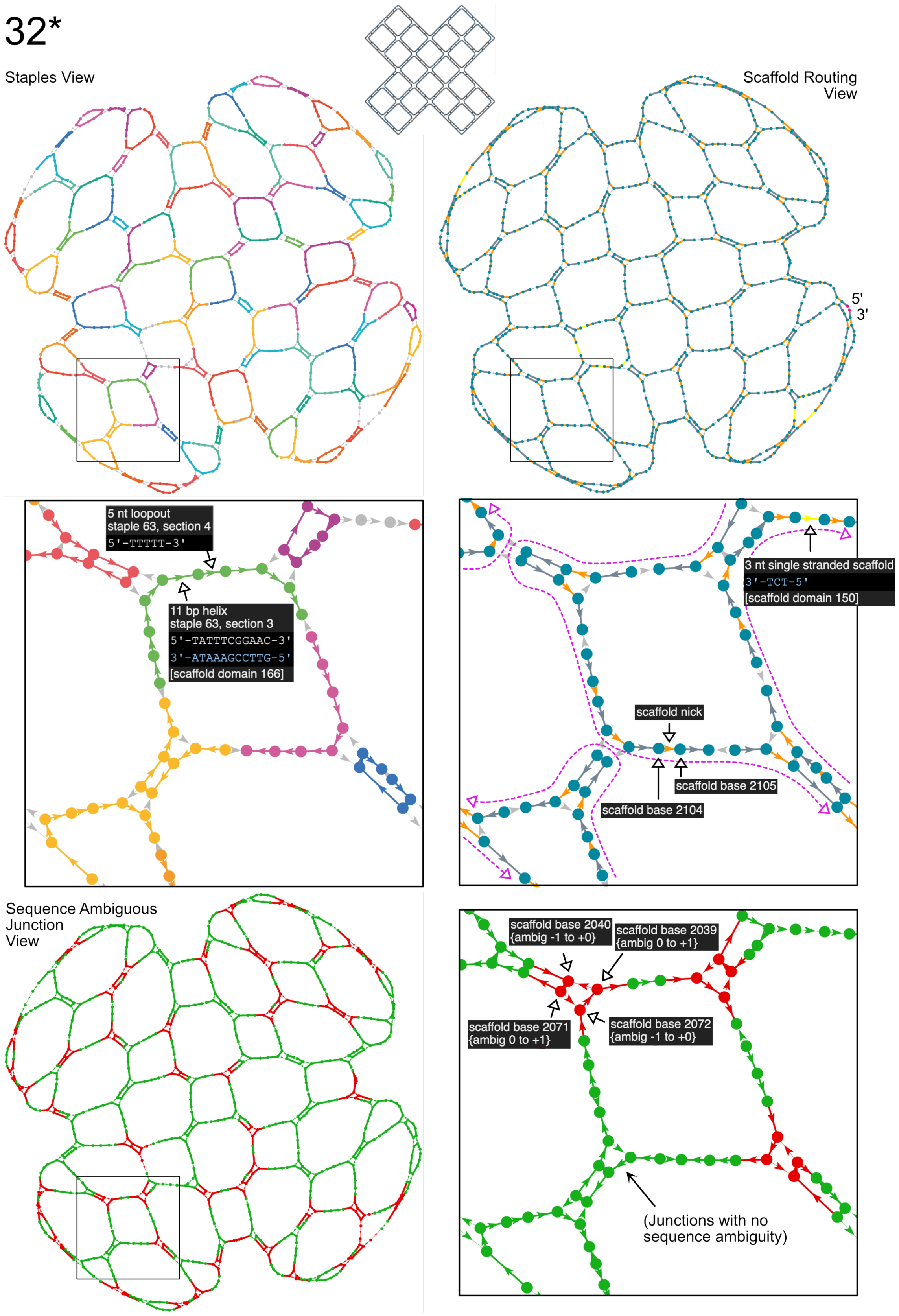


Supplementary Figure 11: Origami 32 (Cross Mesh). Reverse engineered guide schematic shown in three different views and zooms, to highlight features of the interactive HTML interface.

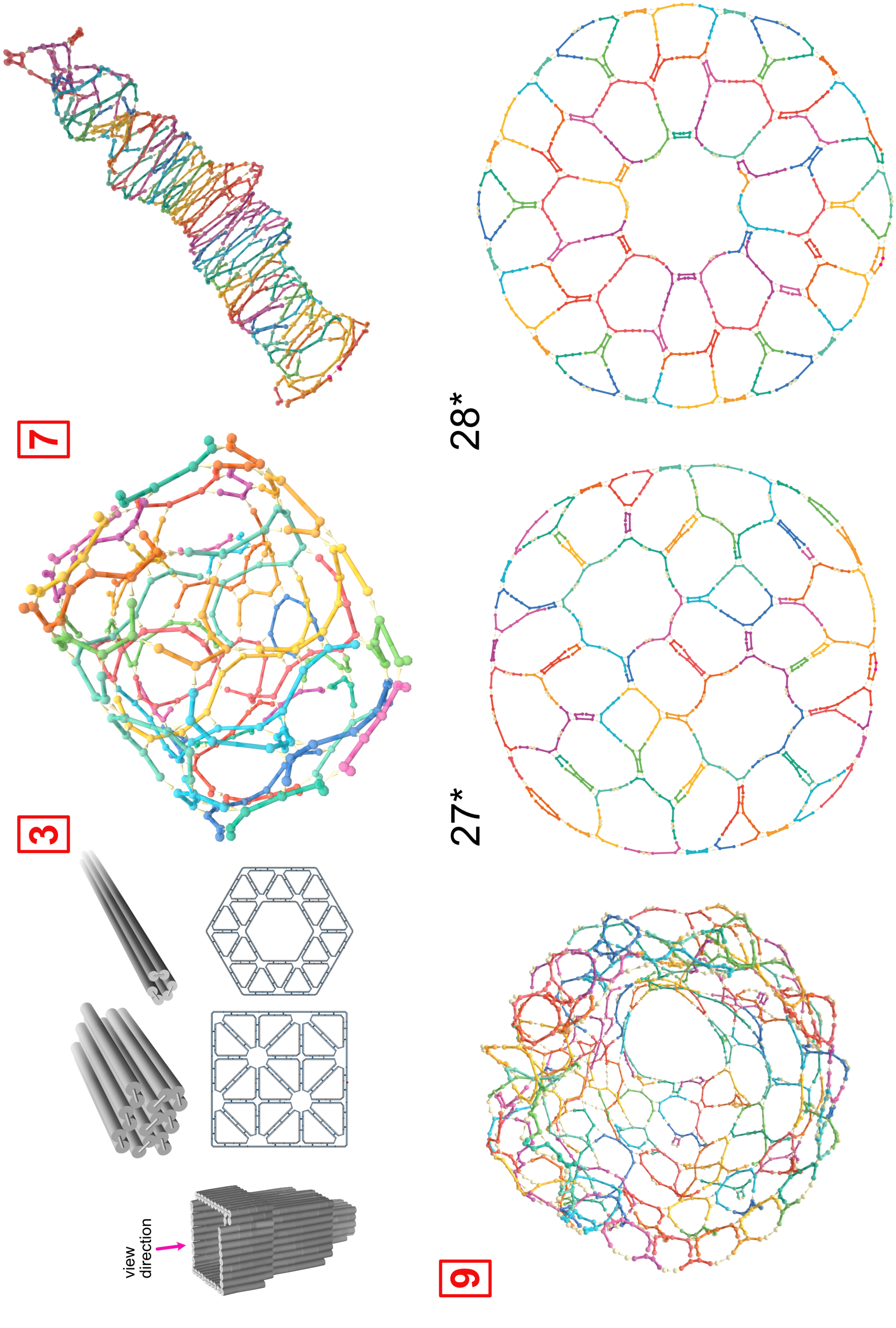


Supplementary Figure 12: Origamis 3, 7, 9, 27, 28 rendered from ground truth contact maps.

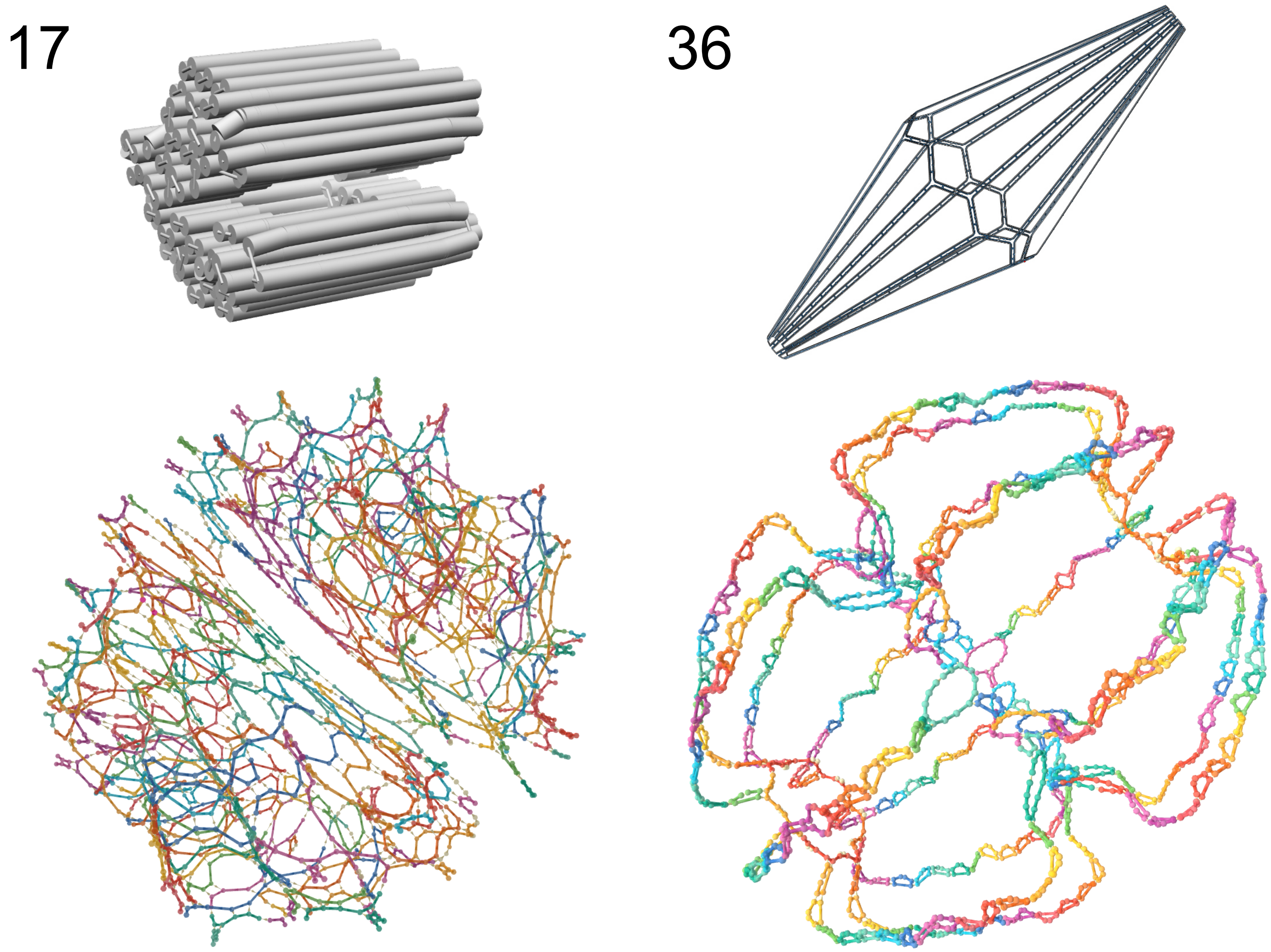


Supplementary Figure 13: Origamis 17 and 36 rendered from ground truth contact maps.
